# A Versatile Antibody Capture System that Drives Precise *In Viv*o Delivery of mRNA loaded Lipid Nanoparticles and Enhances Gene Expression

**DOI:** 10.1101/2024.08.07.607101

**Authors:** Moore Z. Chen, Daniel Yuen, Victoria M. McLeod, Ken W. Yong, Cameron H. Smyth, Bruna Rossi Herling, Thomas. J. Payne, Stewart A. Fabb, Matthew J. Belousoff, Azizah Algarni, Patrick M. Sexton, Christopher J. H. Porter, Colin W. Pouton, Angus P. R. Johnston

## Abstract

Efficient and precise delivery of mRNA is critical to advance mRNA therapies beyond their current use as vaccines. Lipid nanoparticles (LNP) efficiently encapsulate and protect mRNA, but non-specific cellular uptake may lead to off-target delivery and minimal delivery to target cells. Functionalizing LNPs with antibodies enables targeted mRNA delivery, but traditional modification techniques require complex conjugation and purification, which often reduces antibody affinity. Here, we present a simple method for capturing antibodies in their optimal orientation on LNPs, without antibody modification or complex purification. This strategy uses an optimally oriented anti-Fc nanobody on the LNP surface to capture antibodies, resulting in protein expression levels >1000 times higher than non-targeted LNPs and >8 times higher than conventional antibody functionalization techniques. These precisely targeted LNPs showed highly efficient *in vivo* targeting to T cells, with minimal delivery to other immune cells. This approach enables the rapid development of targeted LNPs and has the potential to broaden the use of mRNA therapies.

## MAIN

There is growing interest and an urgent need to develop precise, controlled, and cost-effective systems to deliver therapeutic messenger RNA (mRNA)[1]. The successful deployment of two mRNA-lipid nanoparticle (LNP) vaccines to combat the COVID-19 pandemic has demonstrated the potential of LNPs to effectively deliver large synthetic mRNA payloads *in vivo*[2, 3]. However, the low delivery efficiency and low specificity of current LNP formulations limits their use. Passive targeting of LNPs is generally achieved by manipulating the lipid formulation (e.g., different lipids or composition) to achieve accumulation in desired organs or cells[4]. Different charged lipids will recruit different serum proteins to the LNPs, resulting in altered *in vivo* biodistribution[5–7]. Library screens of hundreds of LNP formulations have increased our ability to control the biodistribution of different LNPs *in vivo*[7], however this approach remains labour-intensive and lacks a clear understanding of the specific cells that are targeted.

Active targeting can be achieved by employing targeting ligands such as antibodies or small molecule receptor ligands[8–11]. Rurik, et al. recently reported CD5-targeted LNPs that effectively produce anti-fibrotic chimeric antigen receptor T cells in mice[12]. The same approach has been adapted to apply to *in vivo* hematopoietic stem cell targeting with a CD117 (c-Kit) targeted LNP, which has shown effective *in vivo* stem cell editing and near-complete correction of hematopoietic sickle cells *in vitro* and successful delivery of pro-apoptotic PUMA (<u>p</u>53 <u>u</u>pregulated <u>m</u>odulator of <u>a</u>poptosis) mRNA *in vivo* via targeted mRNA-LNP[13]. However, while greatly improving the specificity of LNP delivery, conventional techniques for immobilising antibodies onto particles affect the antibody binding affinity and often require complex and time-consuming purification protocols to isolate the targeted particles. The widely used succinimidyl ester (SE) or EDC/NHS conjugation chemistries rely on reactions with primary amines on lysine amino acids. Coupling via these lysine residues results in antibodies that are randomly oriented on the nanoparticle surface (Fig. 1a) and can inactivate the antigen recognition domains. Previous work has shown this attachment strategy produces significantly poorer cell binding compared to antibodies that are conjugated via a specific conjugation site[14, 15]. To enable greater control over the antibody orientation, disulfide bonds in the hinge region of the antibody can be reduced to enable coupling of the reduced cysteines with maleimide groups. While this ensures the antibodies are all oriented in the same direction, the point of attachment is invariant and unlikely to orient the antibody in the optimal orientation. The coupling is also complicated by the need for selective disulfide bond reduction under precise conditions[16, 17].

**Figure 1.**
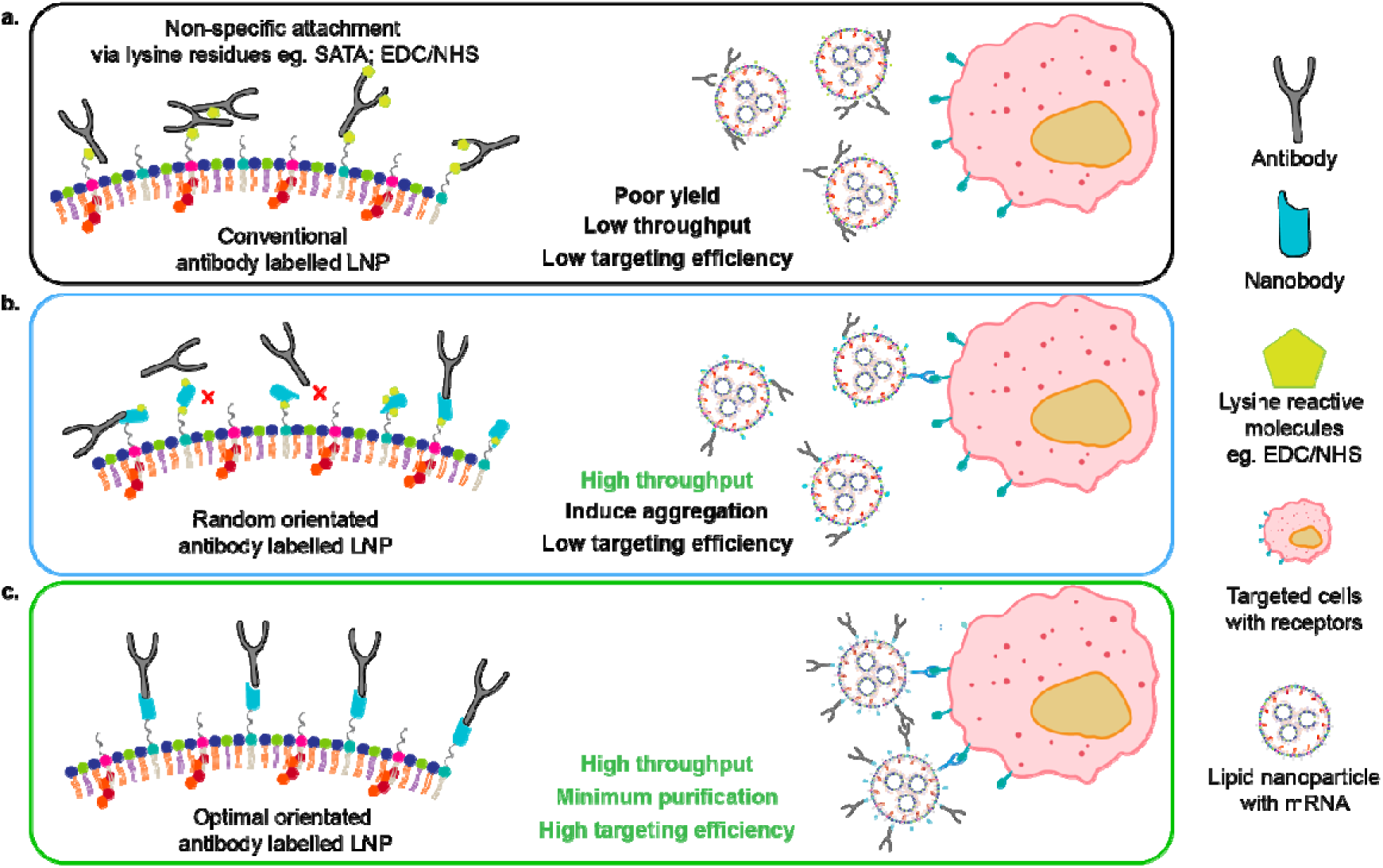
The TP1107 capture system attaches antibodies to LNPs in their optimal orientation. Schematic comparing conventional antibody conjugation methods versus the optimal-orientated antibody capturing system. a) Typically, antibodies are conjugated to nanoparticles by randomly reacting succinimidyl esters with the naturally occurring lysine residues, which compromises the activity of the antibodies as they are randomly oriented on the surface of the nanoparticle. b) Anti-mouse/-rat IgG1 nanobody TP1107 coupled via NHS-Azide to lysine residues allows for high throughput evaluation of antibodies, however, the antibodies are still randomly oriented. c) Our optimally orientated TP1107 captures antibodies in their optimal orientation, maximising binding efficiency and enabling simple and rapid screening of antibodies.

To overcome the limitations of chemical conjugation, non-covalent interactions can be used to attach antibodies onto LNPs. Kedmi et al, reported an innovative and flexible method for antibody-targeted delivery of LNPs (ASSET)[18]. They immobilised an anti-Rat IgG2a single-chain variable fragment (scFv) on to the surface of LNP via peptide lipidation, which served as a modular platform to capture rat IgGa2 antibodies for active cell targeting purposes[8, 19]. While this process eliminated the need to chemically modify antibodies and allowed for efficient and simple coupling to the LNP surface, the N-terminal lipidation of the anti-IgG2a scFv results in sub-optimal antibody orientation on the LNP surface[20]. This leads to less efficient cell targeting and has restricted the ASSET system to capturing a limited range of antibodies.

To overcome this problem, fine control over the protein conjugation site can be achieved through incorporation of synthetic amino acids (SAA), as any residue in a recombinant protein can be substituted using stop-codon reassignment[21]. This approach allows site-specific conjugation for optimal orientation and a well-controlled degree of labelling that will not interfere with the protein structure[22].

Here, we have developed a simple antibody capture system that requires no modification of the antibody, and ensures the antibodies are attached onto LNP in an orientation that increases binding to target by 8 fold compared to conventional antibody capture methods. We have achieved this by modifying the LNPs with a highly specific antibody-capturing nanobody, TP1107[23] (Fig. 1c). To ensure the antibodies are captured on to the surface of LNP with an orientation that maximises binding to the target receptor, we used transmission electron microscopy to determine the position and orientation of TP1107 binding to a mouse IgG1. Using this structure, we identified the optimal position in the TP1107 for conjugation to the LNP surface, and used site-specific incorporation of an azide-bearing synthetic amino acid (p-azido-phenylalanine - azPhe) to allow insertion into the LNP. Due to the high affinity binding of TP1107 to the Fc domain of IgG1, antibodies can be simply added to the TP1107-functionalised LNPs with no requirement for further purification. We demonstrated that optimally oriented antibodies improved LNP binding, and delivery of mRNA more than 1000 times compared with unmodified LNPs, and more than an order of magnitude more than conventional antibody modification. We also demonstrated that this technique can be used to rapidly screen a range of antibodies for optimal specificity and expression efficiency. Using these actively-targeted LNPs we successfully transfected specific cell populations of primary human peripheral blood mononuclear cells (PBMCs) *ex vivo*, with minimal off-target expression. Finally, we demonstrated highly efficient *in vivo* systemic targeting of T cells, with minimal binding to other immune cells. This approach allows for the rapid development of active targeted LNPs and holds the potential to expand the use of mRNA therapies.

## RESULTS

### Rational design and expression of a nanobody to capture antibodies with optimal orientation

Critical to engineering a nanobody that captures antibodies in the optimal orientation is understanding the site where the nanobody binds to the antibody. Previous work established that TP1107 nanobody binds the Fc portion of mouse IgG1[23]. To determine the site of nanobody binding, we used negative stain transmission electron microscopy (TEM) to image the protein complex. A 2:1 ratio of TP1107 and mouse anti-human transferrin receptor (clone OKT9) monoclonal antibody (mAb_TfR_) was negatively stained using uranylformate on a carbon film and imaged at 120 kV. Single particles were picked and extracted using RELION and 2D classification was performed (Supplementary Fig. 1a) [24]. The 2D class averages of the complex reveals a Y-shaped IgG structure along with two additional densities at the terminal ends of the Fc region, corresponding to two nanobodies bound to the antibody (Supplementary Fig. 1b).

The low resolution EM suggests ‘side-on’ binding of TP1107 nanobody to the Fc domain of the IgG structure. This suggests the Gln15 position (highlighted in blue in Supplementary Fig 1c) is a promising site to conjugate TP1107 onto a nanoparticle surface for optimal antibody capture. Therefore, TP1107 with an azido-phenylalanine (AzPhe) at Gln15 (TP1107_optimal_) was produced using a *Methnocaldococcus jannaschii*-derived orthogonal tRNA/synthetase pair[25] and a genomically recoded E.coli host^B-95.ΔA^[26]. We also expressed TP1107 without AzPhe and performed a random lysine modification with NHS-azide (TP1107_random_) (Scheme 1b). A click-reactive fluorescent probe, DBCO-Cy5, was used to determine the efficiency of azide incorporation in these nanobodies. The degree of labeling (DoL) for each batch of TP1107_optimal_ and TP1107_random_ was consistently ∼ 0.5, however due to the random nature of NHS modification, some of the randomly modified nanobodies were modified with multiple azide groups.

### LNP antibody capture system

Next, TP1107_optimal_ and TP1107_random_ were modified with DSPE-PEG_2000_-DBCO lipid (Fig. 2a). The lipid and TP1107 were conjugated at 2:1 molar ratio (DBCO:Azide). 14% of TP1107_optimal_ was coupled to a single DSPE-PEG_2000_-DBCO, as evidenced by the single band at 22kDa using capillary western assay (Fig. 2b). In comparison, 7.9% of the TP1107_random_ was conjugated to a single DSPE-PEG_2000_, with three additional bands, at 26kDa (5%), 32kDa (2.7%) and 36kDa (2.5%) indicating conjugation to 2, 3 and 4 lipids, respectively (Fig. 2b).

**Figure 2.**
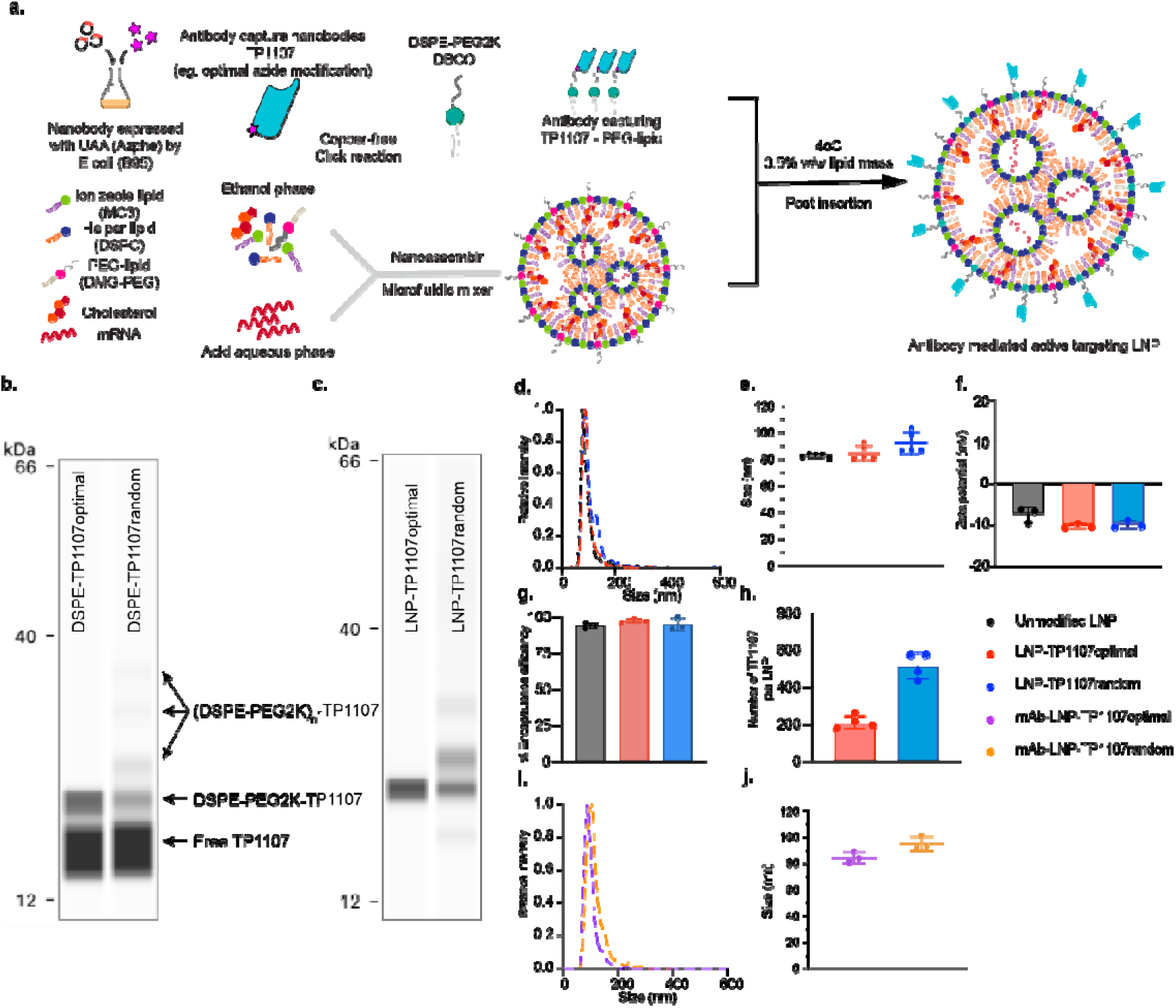
Formulation of LNPs that efficiently capture antibodies in their optimal orientation. Post-insertion of TP1107 into LNPs forms stable particles that efficiently capture antibodies. a) Overview of the process to generate TP1107-LNP. b) ProteinSimple Jess Western assay showing lipid conjugates DBCO-PEG_2000_-DSPE TP1107_optimal_ and DBCO-PEG_2000_-DSPE - TP1107_random_. c) ProteinSimple Jess Western assay showing lipid conjugates inserted into LNP-TP1107_optimal_ and LNP - TP1107_random._ d) Size distribution, e) mode size, f) zeta potential and g) encapsulation efficiency of unmodified LNP, LNP-TP1107_optimal_ and LNP - TP1107_random_. h) Number of TP1107_optimal_ or TP1107_random_ per LNP from 4 different batches of LNP. i) Size distribution and j) mode size of mAb_TfR_ labelled LNP-TP1107_optimal_ and LNP-TP1107_random_. LNPs were synthesized with DLin-MC3-DMA. Data represents mean ± SD (n = 3-5 independent replicates).

LNPs were formulated with clinically approved ionizable lipids (DLin-MC3-DMA or SM-102), using a 50:10:38.5:1.5 molar ratio (ionizable lipid:DSPC:Cholesterol:DMG-PEG_2000_). The hydrodynamic diameter of LNPs measured by NanoSight was 83nm (DLin-MC3-DMA) or 49nm (SM102) (Fig. 2d,e and Supplementary Table S1). To allow nanobody insertion, DBCO-PEG_2000_-DSPE TP1107 mixture was incubated with the LNPs at 0.5% w/w (DBCO-PEG_2000_-DSPE TP1107 vs total lipid) for 48 hours at 4 °C. Unincorporated DSPE-PEG_2000_-DBCO TP1107 was removed by 100kDa MWCO ultrafiltration. Western blotting showed that after purification, no unmodified nanobodies were detected (Fig. 2c). TP1107_optimal_ exhibited a single band, corresponding to nanobody modified with a single DBCO-PEG_2000_-DSPE, while TP1107_random_ exhibited 4 bands corresponding to 1 (57%), 2 (28%), 3 (15%) and >4 (8%) PEG_2000_-DSPE modifications. These multiple bands are consistent with the pattern observed for the DSPE-PEG_2000_-TP1107_random_ (Fig. 2b).

Next, we assessed the basic physical properties of the TP1107 modified LNPs. LNP size was measured using NanoSight, with the diameter of the LNP-TP1107_optimal_ was 85nm and LNP-TP1107_random_ was 92nm, which was similar to the size of the unmodified LNPs (Fig. 2d,e and Supplementary Table S1). We did not expect to see a large increase in the diameter of the LNPs upon functionalization, as the small dimensions of the nanobody (2 x 2 x 4 nm) are within the uncertainty of the NanoSight measurement. LNP-TP1107_random_ also showed a second population sized at 132 nm. This indicated that the randomly labelled TP1107 induced some degree of LNP aggregation. LNP-TP1107_optimal_ showed the same size distribution to the unmodified LNP, with no particle aggregation detected. Attaching TP1107 did not alter the surface charge or the encapsulation efficiency of the LNPs (Fig. 2e,f and Supplementary Table S2). To determine the number of nanobodies attached per particle, the particle concentration was determined using NanoSight (Supplementary Table S3) and the concentration of nanobody was interpolated from calibrated western analysis (Supplementary Table S3). From the ratio of these numbers, the average number of nanobodies per LNP was determined to be ∼ 200 for the LNP-TP1107_optimal_ and 400-600 for the LNP-TP1107_random_ (Fig. 2h). Collectively, these results indicate that the optimally oriented TP1107 is efficiently incorporated into the LNPs without impacting the size of the particles, while the random conjugation method resulted in TP1107 being labelled with more than one DSPE-PEG_2000_-DBCO and caused a small amount of particle aggregation (Fig. 1b).

Functionalization of LNPs with a targeting antibody was achieved by simply adding antibody to the TP1107 functionalized particles at a 2-fold excess of TP1107 to antibody. The high affinity of TP1107 to the Fc domain of mouse or rat IgG1 means under these conditions, all antibody was captured (Supplementary Fig. S2). TP1107_optimal_ LNPs modified with anti-transferrin receptor mouse IgG1 (mAb_TfR_) formed LNPs with a single population with a diameter of 85±4 nm for MC3-LNP and 57±3 nm for SM102-LNP (n=5). The size of the LNPs with randomly captured mAb_TfR_ was similar (95±8 nm for MC3-LNP, n=5). These results confirmed the ability to attach the antibody capturing TP1107 nanobody onto the surface of LNPs, and that this modification did not significantly change the size, surface charge or the ability to efficiently encapsulate mRNA (Supplementary Table S1 & S2). Furthermore, to test whether the nanobody:antibody complex on LNPs is stable in plasma, we incubated the antibody modified LNPs in human plasma at 37°C overnight. We observed no antibody displacement (Supplementary Figure 2). This confirmed robust antibody capture on the surface of the LNPs and that the capture system is suitable as an active targeting strategy for mRNA delivery.

### Optimizing delivery to specific cells

To engineer the most efficient actively targeted LNP system, the base LNP formulation should have minimal non-specific association with cells. For this purpose, we investigated two PEG lipids. First DMG-PEG_2000_, which is used in the Moderna vaccine formulation, and second DSPE-PEG_2000_, which has longer alkyl chains which may anchor the PEG to the surface of the LNP better. Both PEG lipids were formulated with two of the current clinically approved ionizable lipids (SM102, used in the Moderna Vaccine and DLin-MC3-DMA, used in Onpattro). The unmodified DMG-PEG_2000_-LNP or DSPE-PEG_2000_-LNP were formulated with mRNA encoding eGFP spiked with a Cy5 labelled oligo to assess the induced protein expression and association of LNPs to cells, respectively. LNP formulations were incubated with Jurkat cells for 24 hours at 0.5-1 ng/µL mRNA concentration. Untargeted DMG-PEG_2000_-LNPs showed >5-fold more non-specific association compared to untargeted DSPE-PEG_2000_-LNPs after 24 hours (Fig. 3a and Supplementary Fig. 3). For DLin-MC3-DMA, untargeted DSPE-PEG_2000_-LNPs exhibited limited eGFP expression, while untargeted DMG-PEG_2000_-LNP induced strong eGFP expression (Fig. 3b). Similar results were observed with the SM102-LNP formulation (Supplementary Fig. 3). Next, we used our TP1107 capture system to functionalize these LNPs with mAb_TfR_, which binds to human TfR that is expressed on the surface of Jurkat cells. A mouse IgG1 isotype control antibody (mAb_iso_) was used to assess the potential non-specific association induced by modifying the PEGylated LNP surface with proteins. Both mAb_TfR_-DMG-PEG_2000_-LNPs and mAb_TfR_-DSPE-PEG_2000_-LNPs showed significantly increased association with cells compared to unmodified LNPs. The association of both targeted formulations to cells was similar, however the lower non-specific association of the DSPE-PEG_2000_-LNPs resulted in a 61-fold increase in specific binding, compared to an 11-fold increase with DMG-PEG_2000_-LNPs (Fig. 3c). eGFP expression from the mRNA delivered by the LNPs followed a similar trend. mAb_TfR_-DMG-PEG_2000_- and mAb_TfR_-DSPE-PEG_2000_-LNPs both showed high levels of eGFP expression (Fig. 3b). Again, the unmodified and mAb_control_-DSPE-PEG_2000_-LNPs maintained their low non-specificity, with minimal eGFP expression. A >1880-fold increase in eGFP expression was observed with the mAb_TfR_-DSPE-PEG_2000_-LNPs compared to unmodified DSPE-PEG_2000_-LNPs. While the mAb_TfR_-DMG-PEG_2000_-LNPs showed similar eGFP expression to the mAb_TfR_-DSPE-PEG_2000_-LNPs, the DMG LNPs showed a lower 73-fold improvement compared to unmodified LNPs (Fig. 3c) due to the higher non-specific expression of eGFP. We saw similar results for SM102 formulated LNP, (Supplementary Fig. 3). Therefore, from hereon, we formulated all LNPs with DSPE-PEG_2000_ as the PEG component.

**Figure 3.**
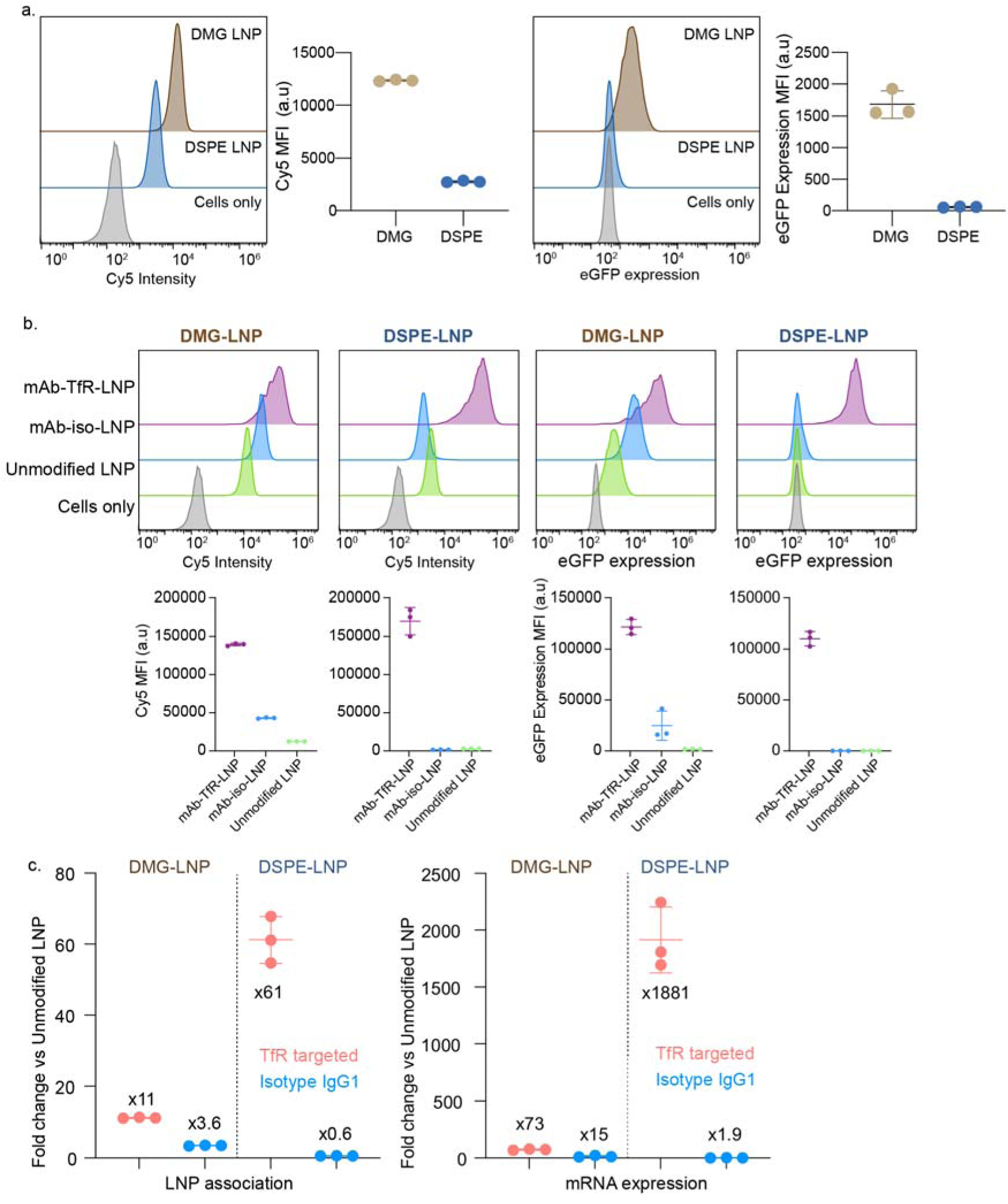
DSPE-PEG_2000_-LNPs limit non-specific cell binding, while maintaining high antibody mediated binding. a) Left panel, Cy5 mean fluorescence intensity (MFI) of Jurkat cells incubated with DMG-PEG_2000_-LNP or DSPE-PEG_2000_-LNP (with DLin-MC3-DMA as the ionizable lipid). Right panel, eGFP expression level of Jurkat cells incubated with DMG-PEG_2000_-LNP or DSPE-PEG_2000_-LNP. b) Cy5 MFI (left) and eGFP expression (right) of Jurkat cells incubated with human TfR targeted LNP, isotype control and unmodified LNP. c) Calculated fold change difference between targeted LNP vs. unmodified LNP for LNP association and mRNA expression in DMG-PEG_2000_-LNP or DSPE-PEG_2000_-LNP treated Jurkat cells. Data represents mean ± SD (n = 3 replicate wells).

To demonstrate the antibody capture system is compatible with different LNP formulations, we also formulated DMG-PEG_2000_-LNP and DSPE-PEG_2000_-LNP with SM102 as the ionizable lipid rather than MC3. SM102-LNP formulations were incubated with Jurkat cells for 24 hours at 0.5 ng/µL mRNA concentration (Supplementary Fig. 3). We observed similar trends with SM102 and MC3-LNPs, with both mAb_TfR_-DMG-PEG_2000_-LNPs and mAb_TfR_-DSPE-PEG_2000_-LNPs (Supplementary Fig. 3a and 3b) inducing significantly higher cell association (Cy5 MFI) and higher mRNA delivery (eGFP expression MFI) compared to the isotype control and unmodified counterpart. Again, DMG-PEG_2000_ showed a higher non-specific association and transfection compared with DSPE-PEG_2000_ (Supplementary Fig. 3c). As has been observed in other studies comparing the effect of the ionizable lipid, SM102 formulations all induced significantly higher protein expression than MC3.

To further tune antibody-capture on LNPs, we investigated how the antibody density on the LNP affected cell binding and the efficiency of protein expression from the delivered mRNA. To determine the efficiency of antibody capture onto the LNPs we used native PAGE followed by the western blot against the antibody to measure the amount of free antibody in solution after functionalization. 100 ng of mAb_TfR_ TP1107_optimal_ or TP1107_random_ LNP with different ratios of antibody:TP1107 (1:64, 1:32, 1:16, 1:8, 1:4, 1:2, 1:1 and 2:1) were loaded into the gel. Free antibody can enter the gel, while LNPs and the antibodies captured on the surface of the LNPs remain trapped in the well. There was no detectable free antibody for 1:64, 1:32, 1:16, 1:8, 1:4, or 1:2 antibody:TP1107 ratios (Supplementary Fig. 4), indicating antibody capture was quantitative, and that these LNPs could be used without any further purification. The 1:1 and 2:1 ratios showed increasing levels of free antibody. To test the cell binding and protein expression induced by the LNPs with different levels of antibody functionalization, we incubated targeted TP1107_random_-LNPs or TP1107_optimal_-LNPs with Jurkat cells for 4 hours without removing unbound antibody. As expected, cell binding (Cy5 fluorescence) and eGFP expression increased as the amount of targeting antibody on the surface of the LNPs increased, with maximum cell association and eGFP expression observed with a 1:8, 1:4 and 1:2 (mAb_TfR_ to TP1107) ratio. At the highest antibody to TP1107 ratios (1:1 and 2:1), the cell binding and eGFP expression significantly decreased (Fig. 5 a-c). This is likely due to free antibody binding to the TfR receptors on the cell surface and blocking the binding of the targeted LNPs. While it is possible to purify the targeted LNPs from the unbound antibodies, this increases the complexity of the functionalization process and will likely result in lower yields of LNPs. From this data we determined the 2:1 TP1107 to antibody ratio was the optimal functionalization ratio, as it requires no further purification of the LNPs, and exhibits significantly higher cell binding and higher protein expression than the unmodified control.

We then investigated whether the optimal orientation of TP1107 (TP1107_optimal_) showed superior cell binding and eGFP expression compared to TP1107_random_ with matching antibody number per LNP (LNP-TP1107_optimal_). LNP-TP1107_optimal_ exhibited ∼ 2 times greater cell binding and > 5 times higher eGFP expression compared to LNP-TP1107_random_ (Fig. 4d-f).

**Figure 4.**
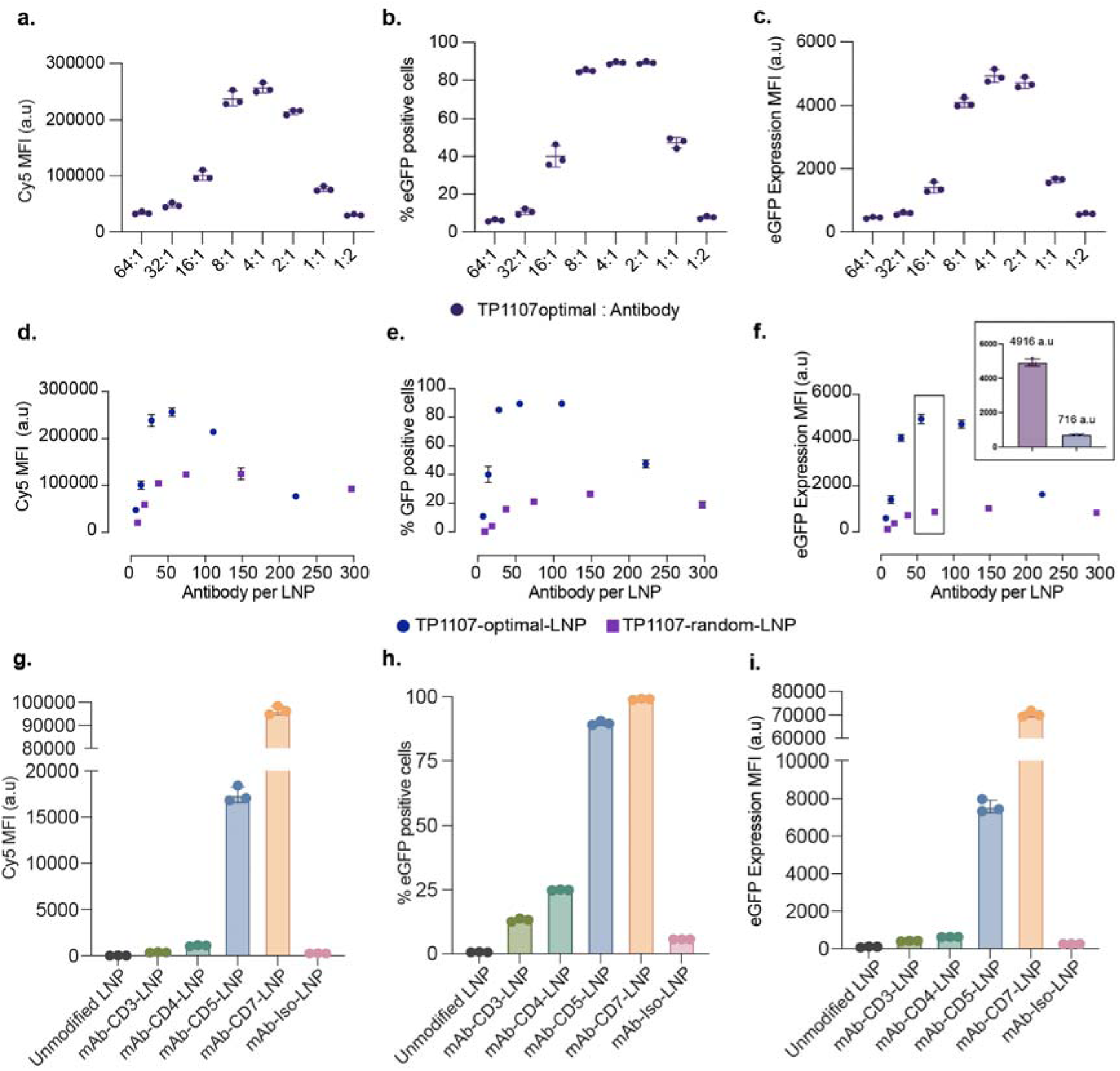
Optimally oriented antibodies show significantly higher cell binding and protein expression than randomly oriented antibodies. a) Cy5 mean fluorescence intensity (MFI), b) percentage of eGFP positive cells and c), eGFP mean fluorescence intensity (MFI) of Jurkat cells incubated with TP1107 optimal incubated with different ratio (64:1 to 1:2) of TP1107 vs mAb_TfR_ over 4 hours with 0.5ng/µL mRNA. Comparison of d) Cy5 mean fluorescence intensity (MFI), e) percentage of eGFP positive cells and f) eGFP mean fluorescence intensity (MFI) of Jurkat cells incubated with TP1107 optimal or random-LNP functionalized with different number of mAb_TfR_ (ranging from 7 to 297 antibody per LNP) over 4 hours at 0.5 ng/µL (with DLin-MC3-DMA as the ionizable lipid). The bar graph represented the data points in the black square. g-i) TP11007_optimal_-LNP functionalized with a panel of mAb (antihuman CD3, CD4, CD5, CD7 and isotype control) were incubated with Jurkat cells for 24 hours. g) Cy5 MFI, h) percentage of eGFP positive cells and i) eGFP MFI. All data are means and error bars represent mean ± SD (n = 3). Data points represent replicate wells. Experiments have been repeated three times.

The strategy currently used to synthesize actively targeted nanoparticles is to directly conjugate a linker to an antibody via reaction to lysine or cystine amino acids. We were interested in comparing this method with our antibody capture system mAb_TfR_ was reacted with NHS-PEG_6_-Azide, which reacts randomly with lysine residues (DOL 0.5). This linker chemistry is similar to the SMCC linker that is also used for antibody/LNP conjugation[12]. Both linkers react randomly with lysine residues, however the SMCC linker contains a maleimide that reacts to thiolated lipids, as opposed to the NHS-PEG_6_-Azide linker, which reacts with DBCO modified lipids. Subsequently, DBCO-PEG_2000_-DSPE lipid was conjugated to the azide modified antibody using the same protocol as the TP1107 lipid conjugation. The LNP and mAb_TfR-Lysine_ mixture was then purified via size exclusion gel chromatography. Then cell binding and mRNA delivery of the mAb_TfR-Lysine_-LNP was compared to the mAb_TfR_-TP1107_optimal_-LNP. While both targeting methods induced 100% eGFP expression in a Jurkat cell line (Supplementary Fig. 5c), the mAb_TfR_-TP1107_optimal_-LNP showed a higher cell association (Supplementary Fig. 5a). More importantly, eGFP expression levels were >8x higher for our optimally oriented antibodies (Supplementary Fig. 5b). These results confirmed that mAb_TfR_-TP1107_optimal_-LNP outperformed the mAb_TfR-Lysine_-LNP.

### TP1107 capture system allows rapid screening of a panel of targeting antibodies

To demonstrate the feasibility of our antibody capture system to rapidly screen a panel of different antibodies, we prepared TP1107_optimal_-LNP with antibodies against human CD3, CD4, CD5, and CD7. These receptors are typically expressed on T lymphocytes, which are targets for drug delivery or chimeric antigen receptor-T (CAR-T) cell therapy[27–29]. First, we measured the surface expression level of these markers on Jurkat cells. There was limited CD3 expression, however there was robust surface expression of CD4, CD5 and CD7 (Supplementary Fig. 6). We next investigated LNP association to the cells. mAb_CD5_ and mAb_CD7_-LNP showed higher levels of binding to Jurkat cells than mAb_CD3_ and mAb_CD4_ (Fig. 4g). Both the mAb_control_ and unmodified LNPs showed similar levels of non-specific association. The association of mAb-LNP to Jurkat cells followed a similar trend to that observed for free mAbs, indicating our antibody-capture method does not interfere with antibody-receptor binding. Interestingly, as was observed for the mAb_TfR_ LNPs, the increase in eGFP expression was even greater than the increase in cellular association for the targeted LNPs. Both mAb_CD5_ and mAb_CD7_-LNPs showed high expression levels (MFI 7584 ± 344 and 70327 ± 1215 respectively) with ∼90-100% of the cells expressing eGFP after 24 hours (Fig. 4h–i). Although the unmodified and isotype LNPs both showed low levels of non-specific association to the Jurkat cells, there was minimal level of eGFP expression (MFI 247 ± 4).

### Targeting specific T cell receptors maximises protein expression in human PBMCs

To further demonstrate the power of this targeted LNP system, we investigated the ability to target specific sub-sets of human peripheral blood mononuclear cells (PBMC). PBMCs were isolated from the blood of healthy donors using a Ficoll gradient purification (Fig. 5a). Human CD3, CD4, CD5 and CD7 targeted LNP with eGFP mRNA and Cy5-labelled oligos were incubated with PBMC for 24 hours. Cells were then analysed by flow cytometry for LNP binding (Cy5 signal) and eGFP expression, using an antibody phenotyping panel for the following population: CD4+ T cells (CD3+, CD4+); CD8+ T cells (CD3+, CD8+); NK Cells (CD3-,CD19-, CD56+); and CD19+ B cells (CD3-, CD19+) (Supplementary Fig. 6a). As expected, CD3, CD4, CD5 and CD7-targeted LNP showed strong binding towards CD4-positive T cells, as these receptors are present on the cell surface. Interestingly, while all CD4+ T cells had targeted LNPs bound (Supplementary Fig. 8a), mAb_CD3_ LNPs showed lower CD4+ T cell association than mAb_CD7_ LNPs, however the eGFP expression was significantly higher (Fig. 5b). Furthermore, while mAb_CD4_ and mAb_CD5_ LNPs showed only slightly lower CD4+ T cell association than mAb_CD3_ and mAb_CD7_ (10-50% reduction in association), the eGFP was significantly less (90% reduction). A similar phenomenon was observed for CD8+ T cells. mAb_CD7_ LNPs showed the highest association with CD8+ T cells, however mAb_CD3_ LNPs showed significantly higher eGFP expression (Fig. 5c). This illustrates the role of the receptor in mediating the uptake and delivery of the mRNA and highlights the importance of screening for the optimal receptor. Only mAb_CD7_ LNPs showed significant binding to NK cells, which resulted in a correspondingly high eGFP signal (Fig. 5d). Both unmodified and mAb_iso_ LNPs showed no detectable binding to T cells, NK cells. As expected, CD19 positive B cells showed no binding nor expression with either unmodified or targeted LNP because of the lack of receptors on the surface (Fig. 5e). To understand the relationship between LNP binding and eGFP expression, we ratioed the eEGFP expression to the Cy5 binding signal for all the samples where the Cy5 signal was significantly higher than the background signal (Fig. 5f). These results confirmed that LNPs targeted to CD3 are significantly more efficient at inducing protein expression. [30–32]

**Figure 5.**
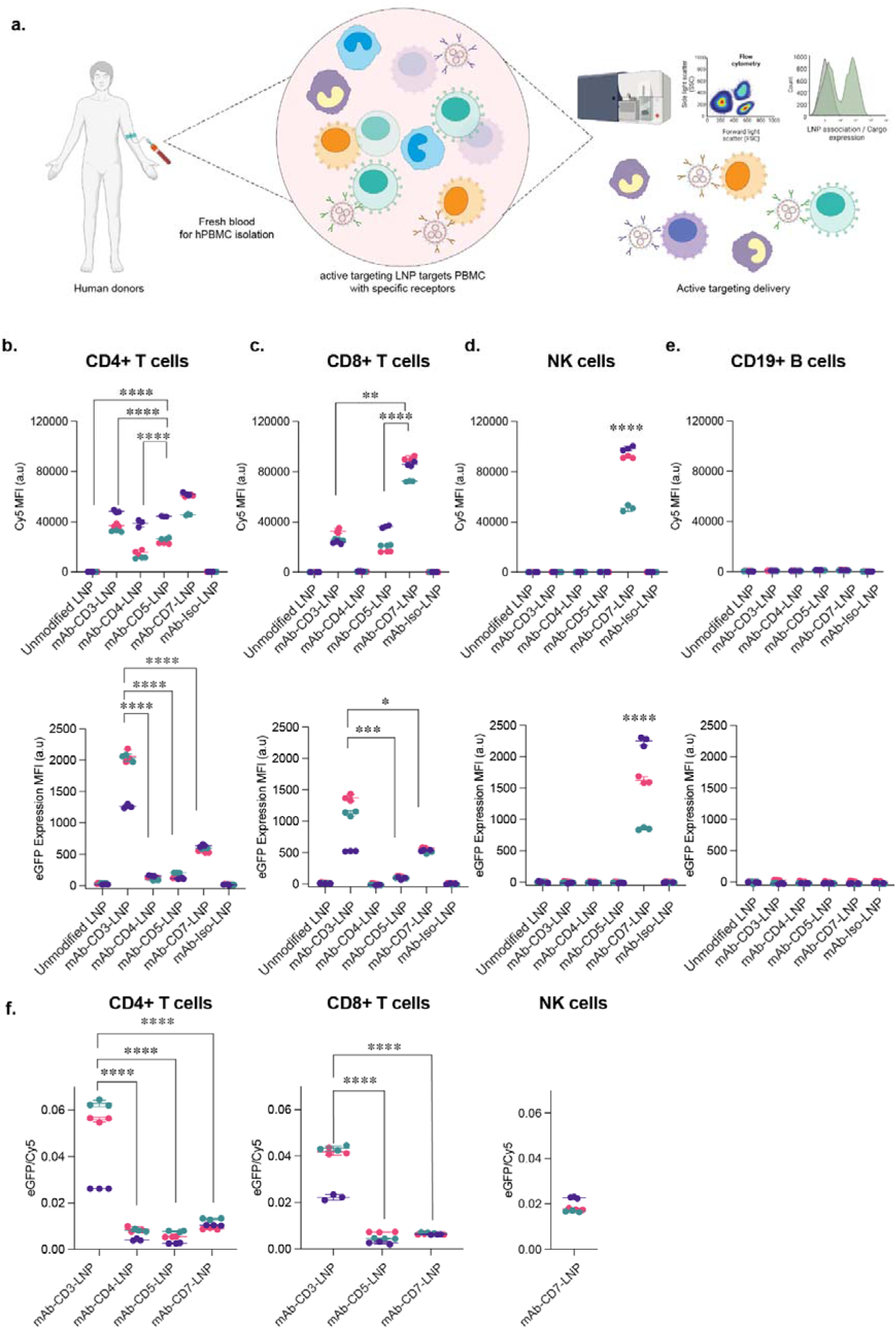
A screen of T cell targeting antibodies in human peripheral blood mononuclear cells (PBMCs) reveals that LNPs targeted to CD3 induces efficient protein expression in T cells. a) Schematic graph illustrates that the process of collecting human PBMC and accessing the specific targeting ability of active targeting LNP system by flow cytometry. b-e) Human PBMC was incubated with a panel of mAb-LNP (CD3, CD4, CD5, CD7, isotype control and unmodified LNP) at 2ng/µL mRNA concentration for 24 hours b) CD4+ T cell, c) CD8+ T cells population, d) CD56+ NK cell population e) CD19+ B cell population. Top panel represented LNP association (Cy5 MFI), bottom panel displayed the mRNA delivery (eGFP MFI). f) Ratio between eGFP MFI/Cy5 of groups that have meaningful LNP association (Cy5 MFI > 3000 and ratio > 0). Data was collected from three donors (depicted by colours green, pink and purple) and conducted in triplicate experiments. Data points represent means of each donor and error bars represent SD. *P < 0.0*5,* \**\**P < 0.0*1,* \**\**P < 0.0*01,* ****P < 0.0001; by two-way ANOVA with post hoc Tukey’s test (Compare row means - main row effects).

We were also interested to test if the superior targeting ability of our system was sustained when using a more potent ionizable lipid. CD3, CD4, CD5, and CD7 targeted LNP as well as IgG1 control and unmodified LNPs, were prepared with eGFP mRNA and SM102 as the ionizable lipid. Mirroring what we observed with the MC3 LNPs, CD3, CD4, CD5, and CD7 targeted LNPs induced a significant increase in the eGFP expression in CD4+ T cells compared to the IgG1 isotype control (Supplementary Figure 9a). In CD8+ T cells, the CD3, CD5, and CD7 but not CD4 targeted LNPs, demonstrated increased eGFP expression (Supplementary Figure 9b). anti-CD7 LNPs targeted both T cells and NK cells, which was also consistent with the CD7 targeted MC3-LNP (Supplementary Figure 9c). No off targeted delivery to CD19+ B cells was observed with any of the antibody targeted LNPs (Supplementary Figure 9d). This indicates that the enhanced expression induced by our targeting system is likely to be applicable to a range of ionizable lipids.

To further demonstrate that our targeted LNP system can target different cell types, we investigated whether we could specifically induce expression in B cells by targeting the LNPs to the CD22 receptor. The CD19 positive B cells population showed a significant increase of eGFP MFI upon the treatment with mAb_CD22_ LNPs, with no expression detected in B cells when PBMCs were treated the isotype control LNPs (Supplementary Figure 10). Importantly, mAb_CD22_ LNPs did not induced eGFP expression in CD22^-^ cell populations (CD4+, CD8+ T cells and NK cells - Supplementary Figure 10). This demonstrates that our targeted LNP system can be engineered to targeted specific cell types depending on the antibody captured on the LNP.

### Targeting circulating T cells in vivo

Finally, to demonstrate that the highly specific T cell targeting we observed *ex-vivo* translates *in vivo*, we administered CD3 targeted LNPs loaded with Cre mRNA into Ai14 mice. These mice have a loxP-flanked STOP cassette preventing transcription of a CAG promoter-driven tdTomato. When Cre recombinase is delivered to any cell in these mice, the STOP cassette is excised and strong tdTomato expression is observed. This allows us to sensitively determine which immune cell population the Cre mRNA is delivered to *in vivo*. 24 hours after i.v. injection of 0.1mg/kg of mRNA (or ∼2µg of mRNA per 20g mouse), the mouse blood was collected via cardiac puncture, the mice liver, spleen and lymph nodes (inguinal, iliac and cervical) were collected and purified to enrich immune cells. The major immune cell populations were phenotyped (Supplementary Figure 7) and percentage of tdTomato cells in each subpopulation was used to quantify successful delivery of Cre mRNA (Figure 6a).

**Figure 6.**
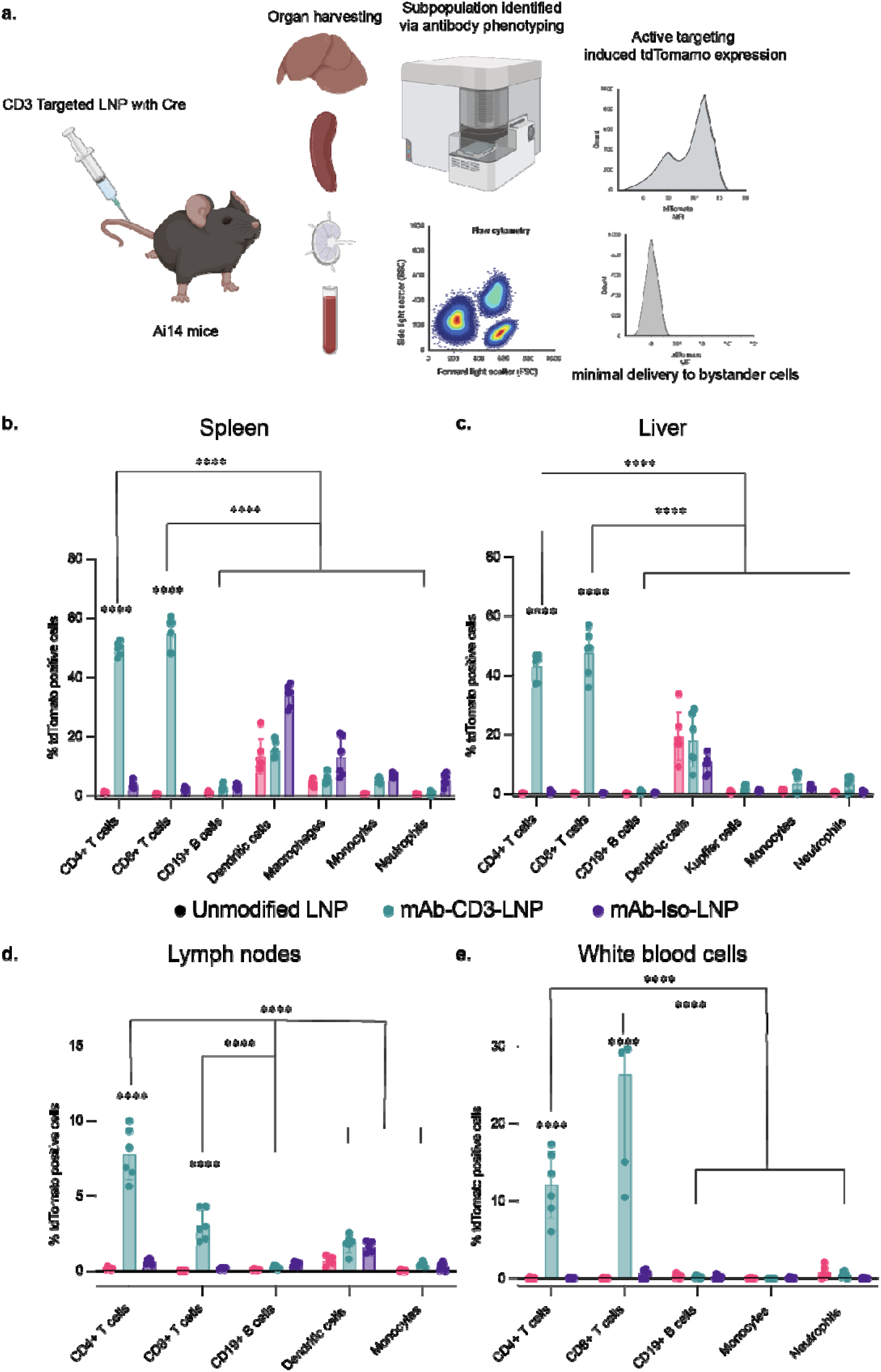
anti-CD3 LNP targeted T cells resident in spleen, liver, lymph nodes and white blood cells *in vivo* with minimal off targeting effect. a) Schematic graph demonstrated that Ai14 mice was given 0.1mg/kg of either CD3 targeted LNP, isotype control LNP and unmodified LNP with Cre mRNA intravenously. Immune cells were harvested from spleen, liver, lymph nodes and circulating blood after 24 hours. b) murine splenocytes, c) murine liver immune cell population, d) murine lymph node immune cell population and e) murine white blood cells, were isolated and enriched. The stained cells were analyzed by flow cytometry to identify the percentage of tdTomato positive cells in each sub population (CD4 + T cell, CD8 + positive T cell, dendritic cells, CD19 + B cells, monocytes, macrophage/kupffer cells, and neutrophils.) n = 6 biologically independent mice in two separate cohorts. All data are means and error bars represent SD. ****P < 0.0001; by two-way ANOVA with post hoc Tukey’s test (Compare row means - “Compare cell means with others in its row and its column”).

We observed that mAb-CD3-LNP induced a significant increase of tdTomato positive T cells population in spleen (50% of CD4+ and 54% CD8+ T cells), liver (43% of CD4+ and 47.5% CD8+ T cells), lymph nodes (8% of CD4+ and 3-4% of CD8+ T cells) and circulating blood (12% of CD4+ and 26% of CD8+ T cells) compared to its isotype control and unmodified LNP (below 4% cross all organs) (Figure 6b,c and d). This result indicates that our targeted system can selectively target the cell population expressing specific receptors *in vivo*. This highly specific targeting of our LNP system demonstrates its potential to expand mRNA delivery to therapeutic targets that are not accessible with passive LNP uptake. More importantly, our targeted LNPs did not increase non-specific delivery in bystander cells compared to its isotype control.

We also assessed the potential toxicity of this targeting system *ex vivo* and *in vivo*. To assess whether the active targeting system changes the cytokine profile in whole human blood, LNPs were incubated with 200uL of whole blood for 24 hours at 2ng/uL concentration. The plasma was then extracted and cytokine (IFNg, IL-1a, IL-1b,IL-6,MCP-1, MIP-1a and TNFa) concentration was measured (Supplementary Figure 11).

All mAb modified LNPs exhibited elevated levels of IL-1b, and TNF compared to unmodified LNPs. This stimulation is likely due to using mouse derived antibody with human cells. mAb-CD3-LNPs also exhibited increased IFNg and IL-1a, although the other mAb modified LNPs exhibited similar levels to unmodified LNPs, suggesting the IFNg and IL-1a is linked to the specific antibody, rather than the LNP formulation. This anti-CD3 mediated activation was anticipated as a number of reports have shown anti-CD3 antibodies can activate T cells via the binding to the CD3ε:TCR complex[33].

For the *in vivo* delivery system, we also found no significant weight loss upon the delivery of mAb-CD3-LNP compared to the current standard delivery formulation and vehicle (Supplementary Figure 12a). Although there was a slight increase in spleen weight in mice treated with mAb-CD3-LNP (Supplementary Figure 12b), histopathological evaluation of the spleen and liver 24 hours post-treatment revealed no remarkable differences (Supplementary Figure 12g). Given the tendency for LNPs to accumulate in the liver, we evaluated the ALT and AST concentrations in plasma at 6 hours post-dose (Supplementary Figures 12c and 12d) and observed no difference between vehicle and the treated groups. A cytokine panel of IL-1a, IL-1b, IL-10, IL-6, MIP-1a, MCP-1, IL-2, TNF•, IFN_γ_, and IL-4 were measured in the plasma at 24 hours post-dosing.

Of the cytokines evaluated, only IL-4 and MCP-1 (CCL-2) were above the limit of detection (10 pg/mL). All four groups (Vehicle, unmodified SM102-DSPE, unmodified SM102-DMG as the current common formulation, and mAb-CD3-LNP) showed no significant difference in IL-4 levels. However, the mAb-CD3-LNP group exhibited a small increase in MCP-1 levels 24 hours post-dosing (Supplementary Figures 12e and 12f). This is consistent with other reports that show elevated levels of MCP-1 (CCL-2) after treatment with anti-CD3 *in vivo*[34].

## DISCUSSION

With increasing interest in using mRNA in therapeutic applications (as opposed to vaccines), there is an urgent need to design a potent, well controlled, and more efficient mRNA delivery system. While the success of the COVID vaccine formulations has demonstrated the potential of mRNA, fulfilling its broader therapeutic potential is likely to require more efficient, and more precisely controlled delivery methods. Active targeting of LNPs allows specific cells to preferentially take up nanoparticles via receptor-mediated endocytosis, and has yielded promising results for the delivery of a range of (non-vaccine antigen) cargos (e.g. CRISPR-Cas9, CAR) to specific cell types[12, 35]. However, current state-of-the-art methods to generate targeted LNPs rely heavily on the non-specific attachment of antibodies via lysine residues or thiol groups. This results in randomly oriented antibodies on the surface of the LNP, which adversely impacts the antibody’s binding to its target antigen due to poor accessibility of the antigen binding domains. It is difficult to control the functionalization process, which often results in multiple conjugation reactions occurring on each antibody, further compromising the activity and potentially inducing aggregation of the LNPs. This approach also requires the isolation of the antibody functionalized LNPs from the free antibody. This purification is challenging due to the high molecular weight of the antibody, and often results in low yields. The complexity of purification limits the number of different targets that can be rapidly explored.

To address these issues, we have developed a highly versatile antibody capture process that attaches antibodies to the surface of LNPs in their optimal orientation. We used an EM model of the nanobody:antibody complex to rationally design the nanobody conjugation site to LNPs so that it was diametrically opposite to the binding interface of the nanobody. From this model we chose the Gln15 position. Site specific coupling was achieved by incorporating azidophenylalanine into the nanobody using stop codon reassignment, and coupling DBCO modified PEG_2000_-DSPE lipid to the azido group. This crude mixture, containing the desired TP1107-PEG_2000_-DSPE conjugate as well as excess unmodified TP1107 and unreacted DBCO-PEG_2000_-DSPE, was added to the mRNA/LNP formulation without any further purification. The TP1107-PEG_2000_-DSPE efficiently inserts into the LNPs, while unmodified TP1107 is easily removed using ultrafiltration system. By exploiting the insertion of the lipidated nanobody into the LNP, the need for the challenging separation of unmodified TP1107 from lipidated TP1107 is circumvented. Targeting antibodies are then quantitatively captured onto the LNPs due to the high binding affinity between the TP1107 nanobody and the invariant Fc region of the antibodies. The quantitative capture means no further purification of the LNPs is required after the addition of the antibody. The simple and straightforward assembly process enables the rapid synthesis of a range of LNPs targeted to different surface receptors. Our antibody capture process displays significant advantages over conventional antibody-LNP systems by virtue of binding orientation, efficiency, and simplicity of manufacture.

For optimal specificity, the LNP’s interaction with cells should be driven solely by the interaction of the antibody with its target receptor. This requires the LNP to have minimal non-specific association with cells. Cullis and co-workers have shown that DMG-PEG_2000_ rapidly dissociates from the LNP in serum, with >50% dissociated in 2 hours[36]. However, by increasing the alkyl chain length, the LNPs can be engineered to retain >90% of the PEG layer over 16 hours. In the non-targeted clinical formulations of LNPs, the dissociation of the DMG-PEG is advantageous, as it can help drive non-specific uptake into (usually liver) cells. However, for targeted delivery systems it is desirable for the stealth PEG layer to remain bound. Our results comparing the non-specific association of DMG-PEG and DSPE-PEG modified LNPs confirm these findings. Thus, the non-targeted 14-C DMG-PEG LNPs showed significantly higher non-specific association with Jurkat cells than the 18-C DSPE-PEG LNPs. In contrast, the cellular association and eGFP expression of both targeted LNPs was similar and therefore the targeting advantage provided by the targeted 18-C DSPE-PEG LNPs was significantly higher. These data highlight the importance of optimizing LNP formulations specifically for targeted delivery, rather than relying on formulations developed for passive delivery since the latter may select for systems with inherently high non-specific binding. Our method also demonstrated promising adaptability across different formulations.

Next, we demonstrated that capturing antibodies in an orientation that allows the Fab domain to freely engage with the target receptors significantly improved association with the target cells and increased subsequent protein expression. Using our method to capture anti-TfR antibodies on LNPs, protein expression increased by eight times compared to LNPs with randomly oriented antibodies. To demonstrate that our capture system offers a ‘mix-and-go’ solution that reduces complexity, cost and significantly increases speed of targeted LNP production, we targeted LNP to four different surface markers (CD3, CD4, CD5, CD7 and CD22) present on human PBMC’s. The different antibodies were successfully captured onto the LNPs and precisely targeted the predicted subsets of PBMCs. The levels of association to each cell type correlated with levels of surface expression of the respective receptors. However interestingly, protein expression did not directly correlate with LNP binding. While mAb_CD3_ LNPs induced the highest levels of protein expression in CD4+ T cells, they showed lower levels of association than mAb_CD7_ LNPs and similar association levels to mAb_CD4_ and mAb_CD5_ LNPs. This highlights the role of the target receptor in the efficiency of LNP delivery and subsequent protein expression. Previous studies have shown that the TCR/CD3 complex is efficiently internalized and recycled[30] in T cells, while uptake of CD4 is limited. The ability to rapidly synthesize a range of LNPs that target different receptors will be essential to maximizing the delivery efficiency of mRNA, and our capture system easily enables the screening of such receptors. To identify the best receptor for mRNA delivery for a specific population, it seems likely that a receptor that is both highly expressed but that also leads to fast turnover is required and that this will vary depending on the different roles that the receptor can have in various cell populations[37, 38].

To demonstrate that our *in vitro* and *ex vivo* results translated to *in vivo* outcomes, we injected a low dose (100 µg/kg) of CD3-targeted LNPs intravenously into Ai14 reporter mice. As observed *in vitro* and *ex vivo*, the CD3-targeted LNPs demonstrated highly efficient mRNA delivery to T cells, with minimal off-target delivery to circulating B cells, monocytes, or neutrophils. The CD3-targeted LNPs did not increase off target delivery to phagocytic cells such as dendritic cells and macrophages compared to unmodified and IgG isotype control LNPs. These results are encouraging for the future development of non-vaccine mRNA therapeutics.

In summary, our antibody capture system offers two primary advantages over existing methods to prepare antibody targeted LNPs. First, the highly efficient capture system allows simple and rapid preparation of LNPs targeted to a range of different cell surface receptors, without modification of the antibodies or complex and inefficient purification. Second, the antibodies are captured in their optimal orientation, improving LNP targeting and subsequent protein expression. Importantly, we have shown that selection of the surface receptor plays a significant role in the efficiency of protein expression. High receptor expression levels and high LNP association does not necessarily correlate to high levels of protein expression, and the ability of the targeted receptor to internalize is likely to be key to utility. We anticipate that as the therapeutic applications for mRNA mature, precise delivery of the mRNA will play an important role in maximizing therapeutic effect and minimizing off target effects. Our simple antibody capture system will enable rapid screening for the optimal receptors to target. We anticipate that the methods outlined here will have broad applicability for the delivery of a broad range of therapeutic nucleic acids.

## Data availability

Data sets generated during the current study are available from the corresponding author on reasonable request.

## Acknowledgements

A.P.R.J. was supported by an NHMRC Career Development Fellowship (GNT1141551) as well as ARC Discovery Projects (DP210103174) and NHMRC Ideas Grant (GNT2011963). This research was also partially funded by the Victoria State Government through funding support from mRNA Victoria for the Victorian mRNA Innovation Hub.

## REFERENCES

1. Qin, S., et al., mRNA-based therapeutics: powerful and versatile tools to combat diseases. Signal Transduct Target Ther, 2022. 7(1): p. 166.

2. Polack, F.P., et al., Safety and Efficacy of the BNT162b2 mRNA Covid-19 Vaccine. N Engl J Med, 2020. 383(27): p. 2603–2615.

3. Baden, L.R., et al., Efficacy and Safety of the mRNA-1273 SARS-CoV-2 Vaccine. N Engl J Med, 2021. 384(5): p. 403–416.

4. Kim, M., et al., Engineered ionizable lipid nanoparticles for targeted delivery of RNA therapeutics into different types of cells in the liver. Science Advances, 2021. 7(9): p. eabf4398.

5. Naidu, G.S., et al., A Combinatorial Library of Lipid Nanoparticles for Cell Type-Specific mRNA Delivery. Adv Sci (Weinh), 2023: p. e2301929.

6. Cheng, Q., et al., Selective organ targeting (SORT) nanoparticles for tissue-specific mRNA delivery and CRISPR-Cas gene editing. Nat Nanotechnol, 2020. 15(4): p. 313–320.

7. Li, B., et al., Combinatorial design of nanoparticles for pulmonary mRNA delivery and genome editing. Nat Biotechnol, 2023.

8. Veiga, N., et al., Cell specific delivery of modified mRNA expressing therapeutic proteins to leukocytes. Nat Commun, 2018. 9(1): p. 4493.

9. Singh, M.S., et al., Therapeutic Gene Silencing Using Targeted Lipid Nanoparticles in Metastatic Ovarian Cancer. Small, 2021. 17(19): p. e2100287.

10. Tarab-Ravski, D., et al., Delivery of Therapeutic RNA to the Bone Marrow in Multiple Myeloma Using CD38-Targeted Lipid Nanoparticles. Adv Sci (Weinh), 2023: p. e2301377.

11. Su, F.Y., et al., In vivo mRNA delivery to virus-specific T cells by light-induced ligand exchange of MHC class I antigen-presenting nanoparticles. Sci Adv, 2022. 8(8): p. eabm7950.

12. Rurik, J.G., et al., CAR T cells produced in vivo to treat cardiac injury. Science, 2022. 375(6576): p. 91-96.

13. Breda, L., et al., In vivo hematopoietic stem cell modification by mRNA delivery. Science, 2023. 381(6656): p. 436-443.

14. Kang, J.H., et al., Improving immunobinding using oriented immobilization of an oxidized antibody. J Chromatogr A, 2007. 1161(1-2): p. 9–14.

15. Vashist, S.K., et al., Effect of antibody immobilization strategies on the analytical performance of a surface plasmon resonance-based immunoassay. Analyst, 2011. 136(21): p. 4431–6.

16. Spicer, C.D. and B.G. Davis, Selective chemical protein modification. Nat Commun, 2014. 5: p. 4740.

17. Nuhn, L., et al., Targeting protumoral tumor-associated macrophages with nanobody-functionalized nanogels through strain promoted azide alkyne cycloaddition ligation. Bioconjugate Chemistry, 2018. 29(7): p. 2394–2405.

18. Kedmi, R., et al., A modular platform for targeted RNAi therapeutics. Nat Nanotechnol, 2018. 13(3): p. 214–219.

19. Veiga, N., et al., Leukocyte-specific siRNA delivery revealing IRF8 as a potential anti-inflammatory target. Journal of Controlled Release, 2019. 313: p. 33–41.

20. Dammes, N., et al., Conformation-sensitive targeting of lipid nanoparticles for RNA therapeutics. Nat Nanotechnol, 2021. 16(9): p. 1030–1038.

21. Budisa, N., Prolegomena to future experimental efforts on genetic code engineering by expanding its amino acid repertoire. Angewandte Chemie International Edition, 2004. 43(47): p. 6426–6463.

22. Yong, K.W., et al., Pointing in the Right Direction: Controlling the Orientation of Proteins on Nanoparticles Improves Targeting Efficiency. Nano Lett, 2019. 19(3): p. 1827–1831.

23. Pleiner, T., M. Bates, and D. Gorlich, A toolbox of anti-mouse and anti-rabbit IgG secondary nanobodies. J Cell Biol, 2018. 217(3): p. 1143–1154.

24. Zivanov, J., et al., New tools for automated high-resolution cryo-EM structure determination in RELION-3. elife, 2018. **7**: p. e42166.

25. Chin, J.W., et al., Addition of p-Azido-l-phenylalanine to the Genetic Code of Escherichia c oli. Journal of the American Chemical Society, 2002. 124(31): p. 9026–9027.

26. Mukai, T., et al., Highly reproductive Escherichia coli cells with no specific assignment to the UAG codon. Scientific Reports, 2015. 5(1): p. 9699.

27. Zhang, Y., et al., Allogenic and autologous anti-CD7 CAR-T cell therapies in relapsed or refractory T-cell malignancies. Blood Cancer Journal, 2023. 13(1): p. 61.

28. Hu, Y., et al., Genetically modified CD7-targeting allogeneic CAR-T cell therapy with enhanced efficacy for relapsed/refractory CD7-positive hematological malignancies: a phase I clinical study. Cell Res, 2022. 32(11): p. 995–1007.

29. Dai, Z., et al., T cells expressing CD5/CD7 bispecific chimeric antigen receptors with fully human heavy-chain-only domains mitigate tumor antigen escape. Signal transduction and targeted therapy, 2022. 7(1): p. 85.

30. Alcover, A. and B. Alarcon, Internalization and intracellular fate of TCR-CD3 complexes. Critical Reviews™ in Immunology, 2000. 20(4).

31. Rennick, J.J., A.P.R. Johnston, and R.G. Parton, Key principles and methods for studying the endocytosis of biological and nanoparticle therapeutics. Nat Nanotechnol, 2021. 16(3): p. 266–276.

32. Akinc, A., et al., A combinatorial library of lipid-like materials for delivery of RNAi therapeutics. Nat Biotechnol, 2008. 26(5): p. 561–9.

33. Menon, A.P., et al., Modulating T Cell Responses by Targeting CD3. Cancers (Basel), 2023. 15(4).

34. Martin, G.H., et al., Myeloid and dendritic cells enhance therapeutics-induced cytokine release syndrome features in humanized BRGSF-HIS preclinical model. Front Immunol, 2024. 15: p. 1357716.

35. Rosenblum, D., et al., CRISPR-Cas9 genome editing using targeted lipid nanoparticles for cancer therapy. Science advances, 2020. 6(47): p. eabc9450.

36. Chen, S., et al., Influence of particle size on the in vivo potency of lipid nanoparticle formulations of siRNA. J Control Release, 2016. 235: p. 236–244.

37. Billingsley, M.M., et al., Ionizable lipid nanoparticle-mediated mRNA delivery for human CAR T cell engineering. Nano letters, 2020. 20(3): p. 1578–1589.

38. Billingsley, M.M., et al., Orthogonal design of experiments for optimization of lipid nanoparticles for mRNA engineering of CAR T cells. Nano Letters, 2021. 22(1): p. 533–542.

