## Supplementary Fig. 1a for "A Versatile Antibody Capture System that Drives Precise *In Viv*o Delivery of mRNA loaded Lipid Nanoparticles and Enhances Gene Expression"

**Supplementary Figures**

| The summarized data of the size (nm) of formulated particles | | | |
| --- | --- | --- | --- |
| LNP | Size (nm) | SD | n |
| MC3-LNP | 83 | 1.5 | 5 |
| TP1107_optimal_ -MC3-LNP | 85 | 5.4 | 5 |
| TP1107_random_- MC3-LNP | 92 | 8.3 | 5 |
| Ab_optimal_ - MC3-LNP | 85 | 4 | 3 |
| Ab_random_ - MC3-LNP | 95 | 5.2 | 3 |
| SM102-LNP | 49 | 2.2 | 5 |
| TP1107_optimal_-SM102-LNP | 62 | 3.5 | 5 |
| Ab_optimal_ -SM102-LNP | 57 | 2.9 | 3 |

**Supplementary Table 1.** **Summary of the hydrodynamic diameter of different LNPs measured by NTA.** All data are mean ± SD. N represents independent repeats.

| The summarized data of the encapsulation efficiency of formulated particles | | | |
| --- | --- | --- | --- |
| LNP | Encapsulation Efficiency | SD | n |
| MC3-LNP | 94 | 2 | 3 |
| TP1107_optimal_ -MC3-LNP | 97.33 | 1.528 | 3 |
| TP1107_random_- MC3-LNP | 95 | 4.359 | 3 |
| SM102-LNP | 95.50 | 0.88 | 3 |
| TP1107_optimal_-SM102-LNP | 92.73 | 1.31 | 3 |

**Supplementary Table 2.** **Summary of the encapsulation efficiency of different LNP formulations measured by Ribogreen assay.** All data are mean ± SD. N represents independent repeats.

| The quantification of TP1107optimal and TP1107random per LNP | | | |
| --- | --- | --- | --- |
| TP1107 functionalized LNP | Concentration of TP1107  ug/mL | Particle number per mL | sdAb per LNP |
| TP1107_optimal_ #1 | 6.09 | 8.82E+11 | 262 |
| TP1107_optimal_ #2 | 4.09 | 7.62E+11 | 179 |
| TP1107_optimal_ #3 | 5.58 | 9.51E+11 | 196 |
| TP1107_optimal_ #4 | 4.61 | 7.18E+11 | 215 |
| TP1107_random_ #1 | 5.28 | 3.06E+11 | 578 |
| TP1107_random_ #2 | 4.89 | 3.39E+11 | 483 |
| TP1107_random_ #3 | 2.6 | 8.70E+10 | 417 |
| TP1107_random_ #4 | 3.6 | 1.38E+11 | 577 |

**Supplementary Table 3. Summary of the particle concentration and TP1107 concentration for 4 independent LNP formulations.**

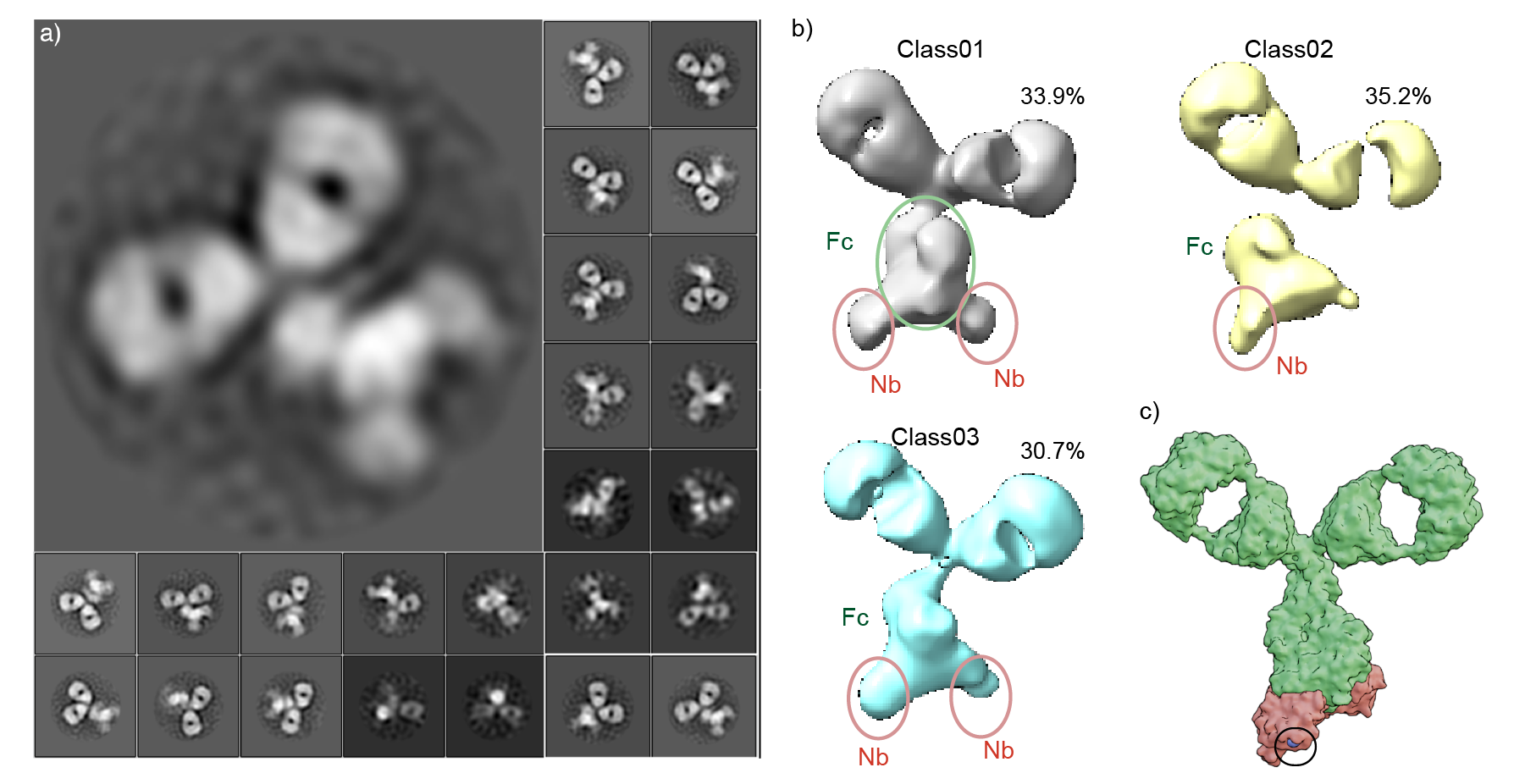

**Supplementary Figure 1. Low resolution EM structure determination enables rational engineering of a nanobody to capture antibodies.** a) TEM obtained for a, 2D projection of nanobody:antibody complexes. Scale bar is 200 Å. b) 3D models showing the nanobody binding to the Fc domain. c) 3D reconstruction of 2D projection from an initial rigid body fitting followed by docking of the nanobodies using HADDOCK. The Gln15 residue (highlighted in Blue and circled in black) was identified as a point of attachment to the LNP that would likely optimize the orientation of the subsequently captured antibody.

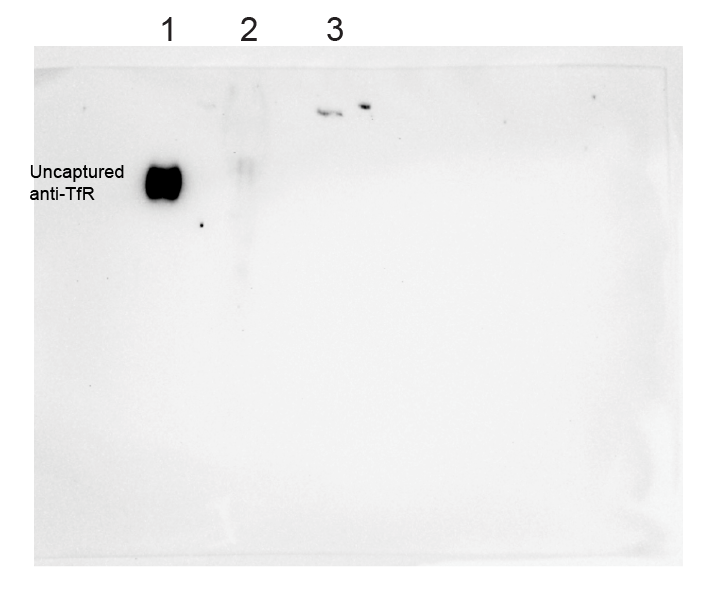

**Supplementary Figure 2. LNP-TP1107_optimal_ with anti-TfR antibody complex remained stable in human plasma.** From left to right, sample loaded as following: Lane 1: 100ng of anti-TfR antibody, Lane 2: 100ng of anti-TfR captured on LNP-TP1107optimal and incubated with human plasma, Lane 3: 100ng of anti-TfR captured on LNP-TP1107optimal and incubated with PBS. The incubation period was 24 hours at 37oC. Protein gel was run under native conditions and followed by western blot detecting mouse antihuman TfR antibody. Only free anti-TfR can enter the gel, when anti-TfR is captured on the LNP it remains in the well. The absence of a band indicates there is no free anti-TfR in solution.

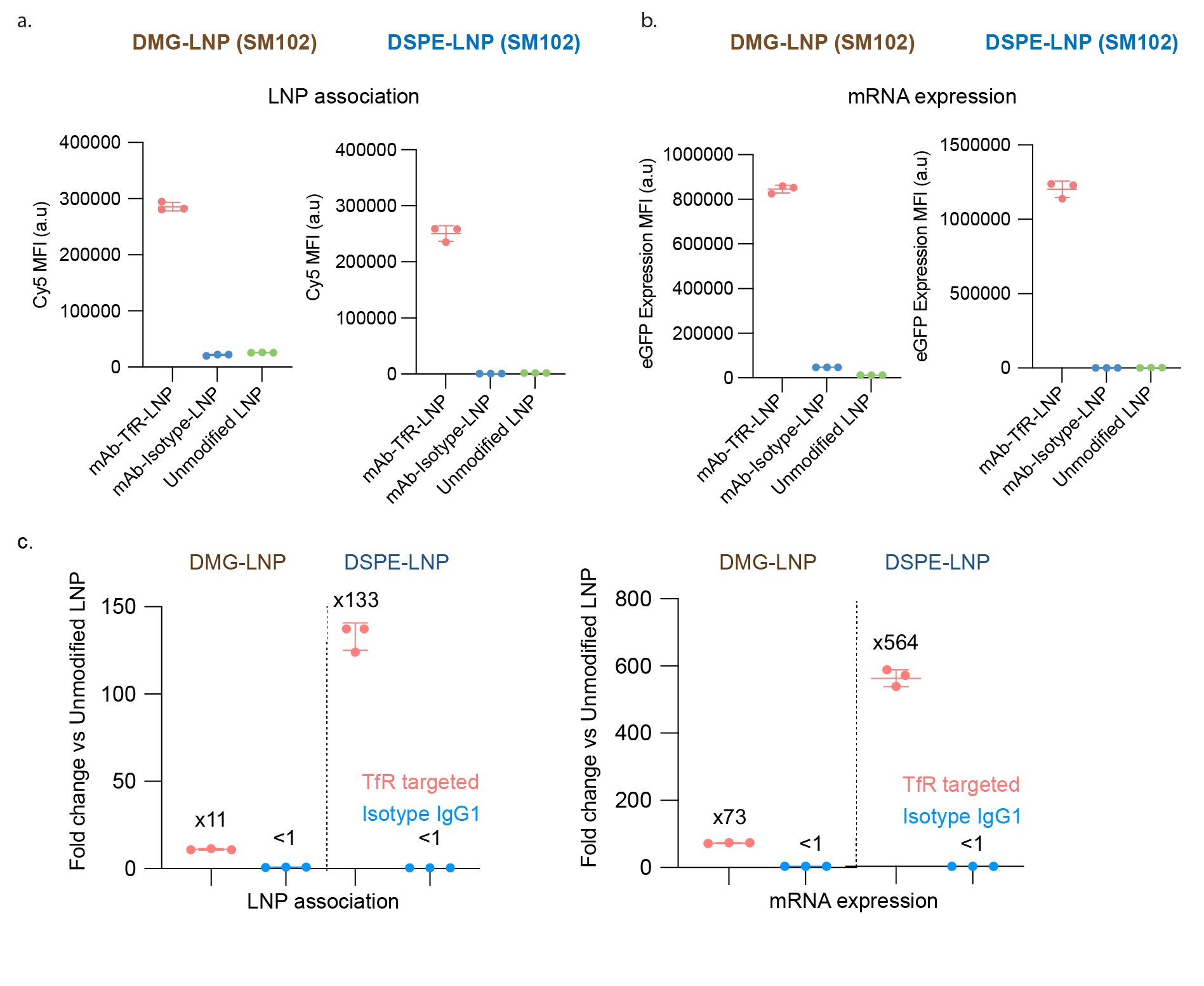

**Supplementary Figure 3. TP1107-PEG2000-DSPE module with SM102-LNP showed enhanced cell binding and mRNA delivery with antiCD71** a) Left panel, Cy5 mean fluorescence intensity (MFI) of Jurkat cells incubated with DMG-PEG2000-SM102-LNP or DSPE-PEG2000-SM102-LNP with either human TfR targeted LNP, isotype control and unmodified LNP. b) eGFP expression level of Jurkat cells incubated with DMG-PEG2000-SM102-LNP or DSPE-PEG2000- SM102-LNP with either human TfR targeted LNP, isotype control and unmodified LNP. Data represents mean ± SD (n = 3 replicate wells).

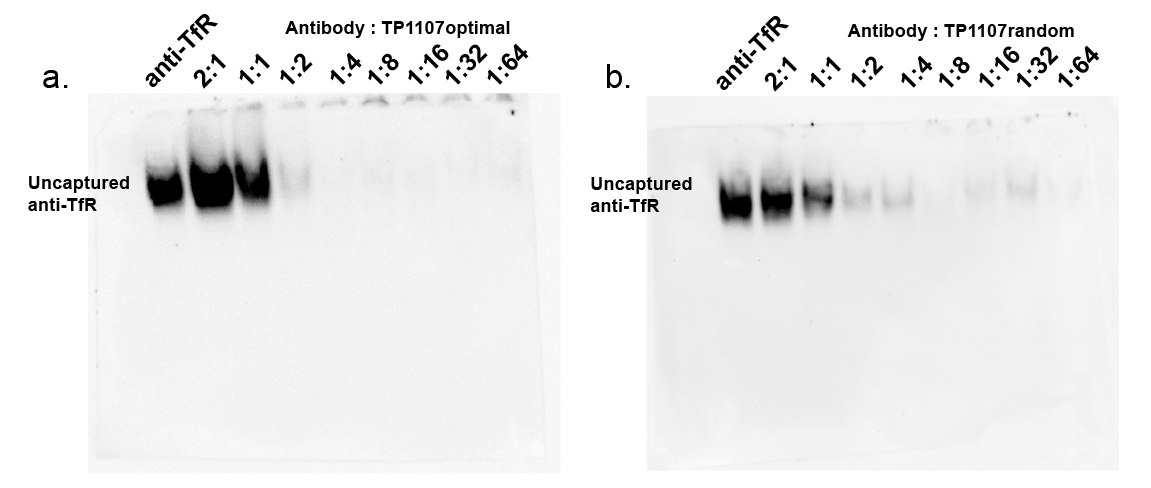

**Supplementary Figure 4. LNP-TP1107_optimal_ successfully captured all antibodies in solution with 1:2 antibody vs TP1107_optimal_ or lower ratio.** From left to right, sample loaded as following: 100ng anti-TfR antibody, 2:1, 1:1, 1:2, 1:4, 1:8, 1:16, 1:32, and 1:64 of antibody (100ng) vs TP1107optimal or TP1107random. Protein gel was run under native conditions and followed by western blot detecting mouse antihuman TfR antibody.

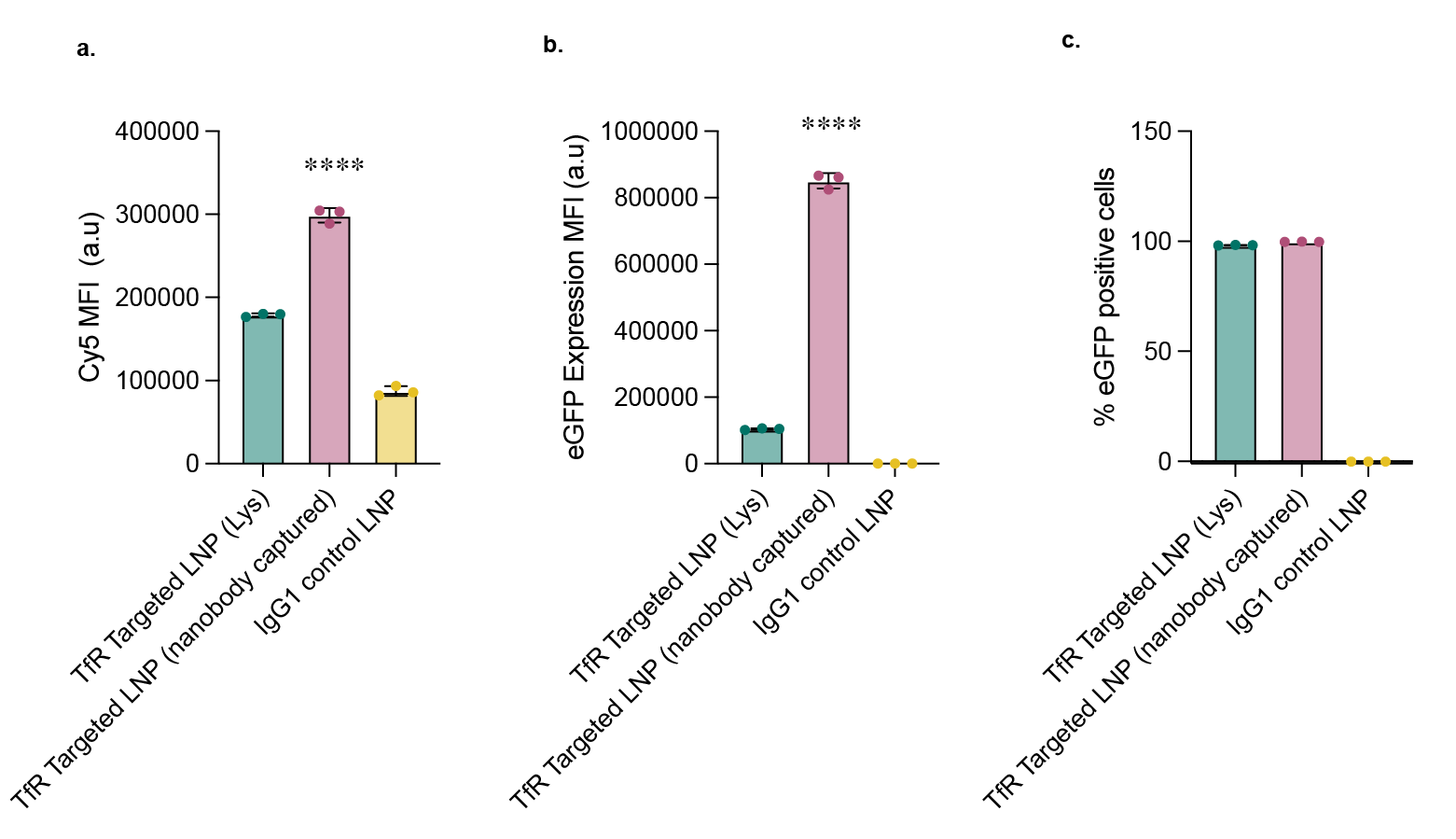

**Supplementary Figure 5. TP1107 captured antibodies outperformed the conventional labelled antibodies on LNP over 24 hours**. a) Cy5 MFI, b) eGFP MFI and c) percentage of eGFP positive cells, of Jurkat cells incubated with mAbTfR-LNP-Lysine reacted or mAbTfR-LNP-TP1107optimal for 24 hours at 0.5 ng/µL mRNA concentration. P value calculated by one-way ANOVA with post hoc Tukey’s test. ****P<0.0001. All data represented as mean ± SD (n = 3).

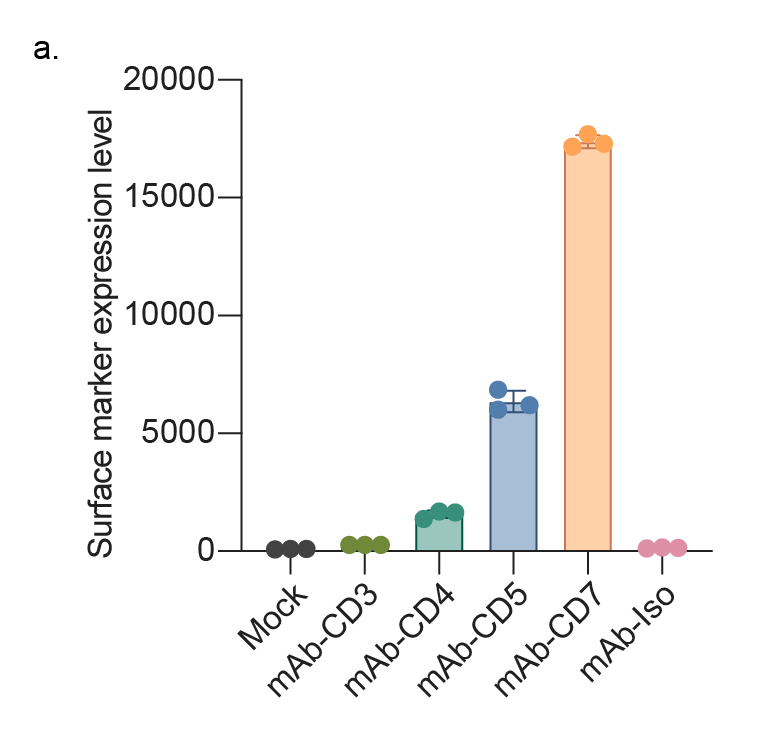

**Supplementary Figure 6. The expression level of CD2, CD3, CD4, CD5 and CD7 on Jurkat cells.** Jurkat cells were incubated with antibodies at 1:100 dilution for 1 hour at 4^o^C. Cells were washed with cold PBS three times before staining with goat anti-mouse Alexa Fluor 647 at 1:1000 dilution for 1 hour at 4^o^C. The mean fluorescent intensity (MFI) of Cy5 was collected and analyzed as surface marker expression level. Mock sample representative cells were only incubated with the secondary antibody. All data are represented as mean ± SD (n = 3).

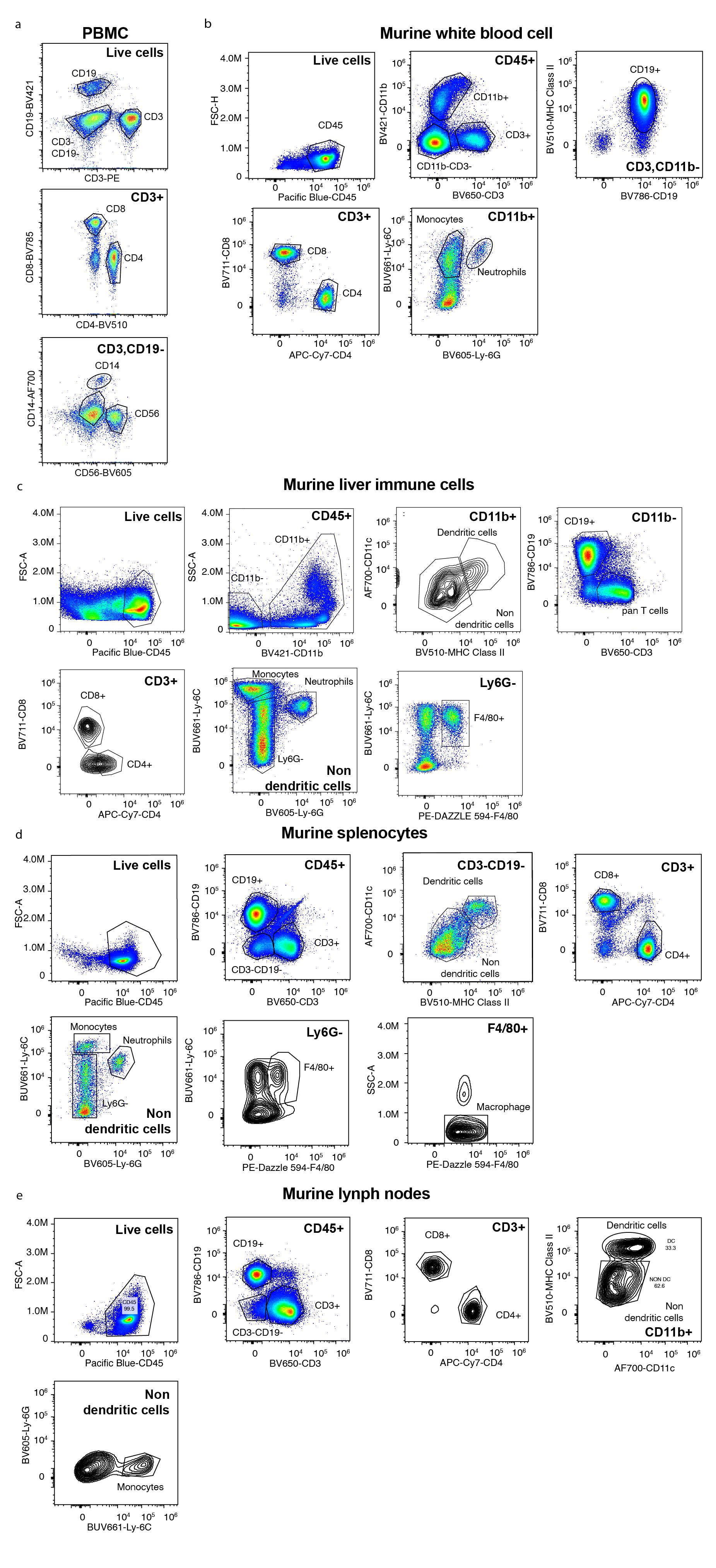

**Supplementary Figure 7. Gating strategy of identifying subpopulation in purified human PBMC, murine white blood cells, murine splenocytes, murine liver immune cell population, and murine lymph node immune cell population. a)** Purified human PBMC were stained with an antibody cocktail (Live and Dead dye, antiCD3, antiCD19, antiCD4, antiCD8a, antiCD14 and antiCD56). Live cells were selected first then subphenotyped into different groups as shown in the figure. **b)** enriched mice WBC, c), enriched murine liver immune cells d) enriched murine splenocytes immune cells and e) enriched murine lymph nodes immune cells were stained with an antibody cocktail (Live and Dead dye, antiCD45, antiCD3/antiCD90.2, antiCD19, antiCD4, antiCD8a, antiMHCII, antiCD11c, antiF4/80, antiLy6C and antiLy6G). The cell population was selected based on FSC-A vs SSC-A and followed by selecting single cells based on FSC-A vs FSC-H. Live cells were selected as the negative population of the live/dead dye.

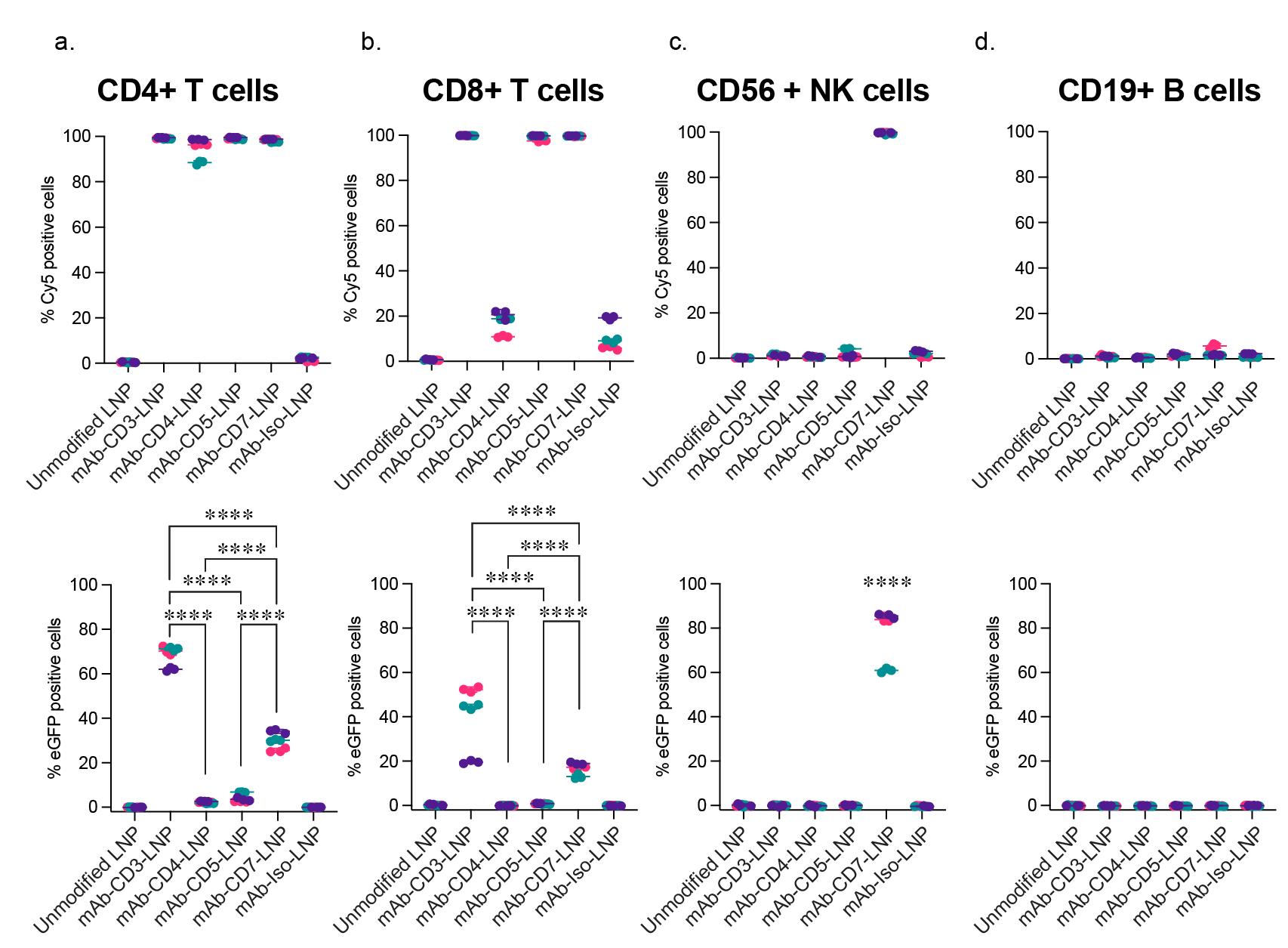

**Supplementary Figure 8. The plotting of percentage of LNP association cells (Cy5 positive population,) and the percentage of EGFP expressing cells (eGFP positive population) with different immune cell population that were treated with mAb-LNP.** The dataset is an extension of Figure 6. a, CD4+ T cell, b, CD8+ T cells population, c, CD56+ NK cell population d, CD19+ B cell population. Cluster dot represents individual donors (depicted by colours green, pink and purple). ****P < 0.0001; two-way ANOVA and Tukey’s post-test (Compare row means - main row effects). All data are means ± SD; n = 3 independent donors.

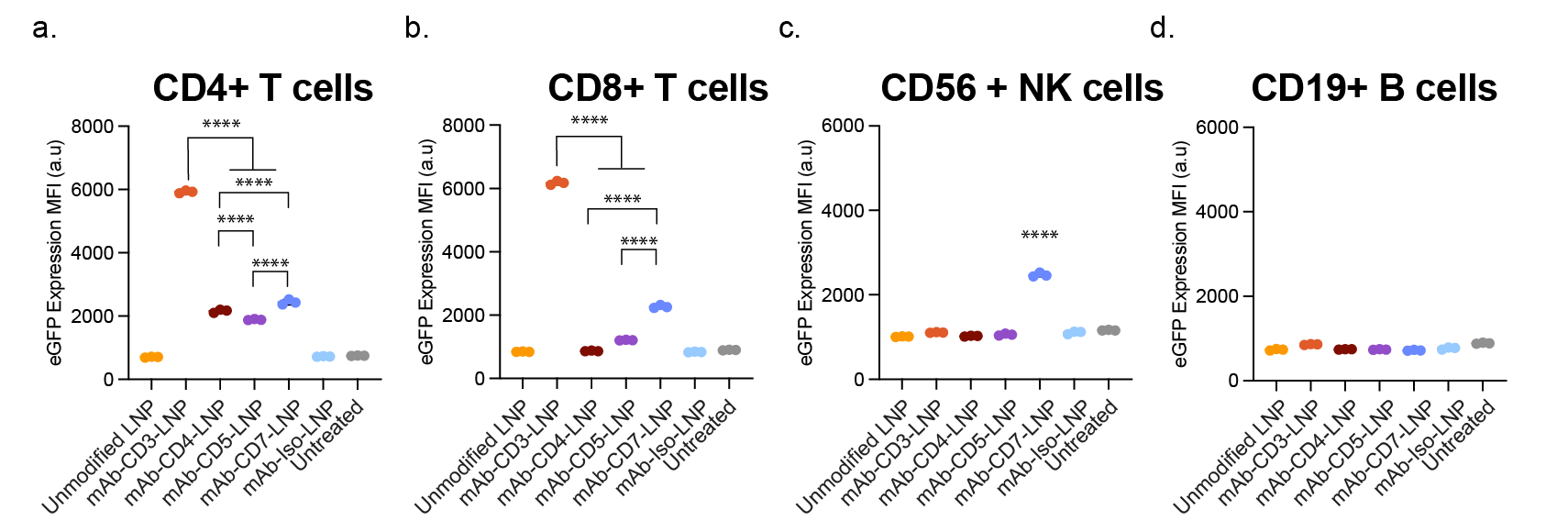

**Supplementary Figure 9. The eGFP expression MFI of different immune population in human PBMC treated with different targeted LNP with SM102 as the ionizable lipid**. a, CD4+ T cell, b, CD8+ T cells population, c, CD56+ NK cell population d, CD19+ B cell population. ****P < 0.0001; one-way ANOVA and Tukey’s post-test. All data are means ± SD; n = 3 independent wells.

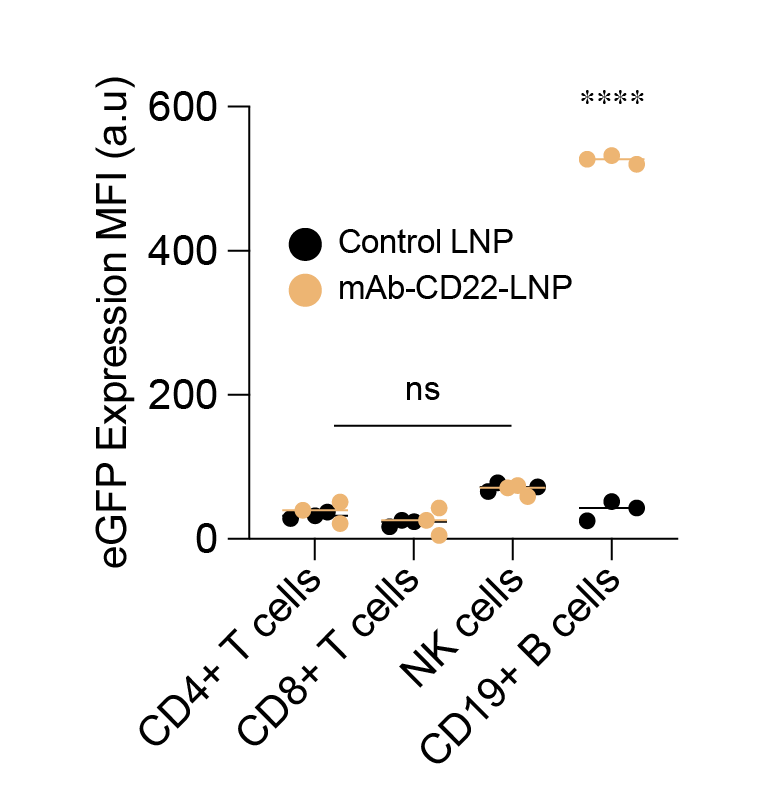

**Supplementary Figure 10. The eGFP expression MFI of different immune population in human PBMC treated with CD22 targeted LNP.** human PBMC cells were purified from fresh donated blood and incubated with either CD22 targeted LNP or the isotype control at 2ng/uL for 24 hours. eGFP expression was plotted. ****P < 0.0001; one-way ANOVA and Tukey’s post-test. All data are means ± SD; n = 3 independent wells.

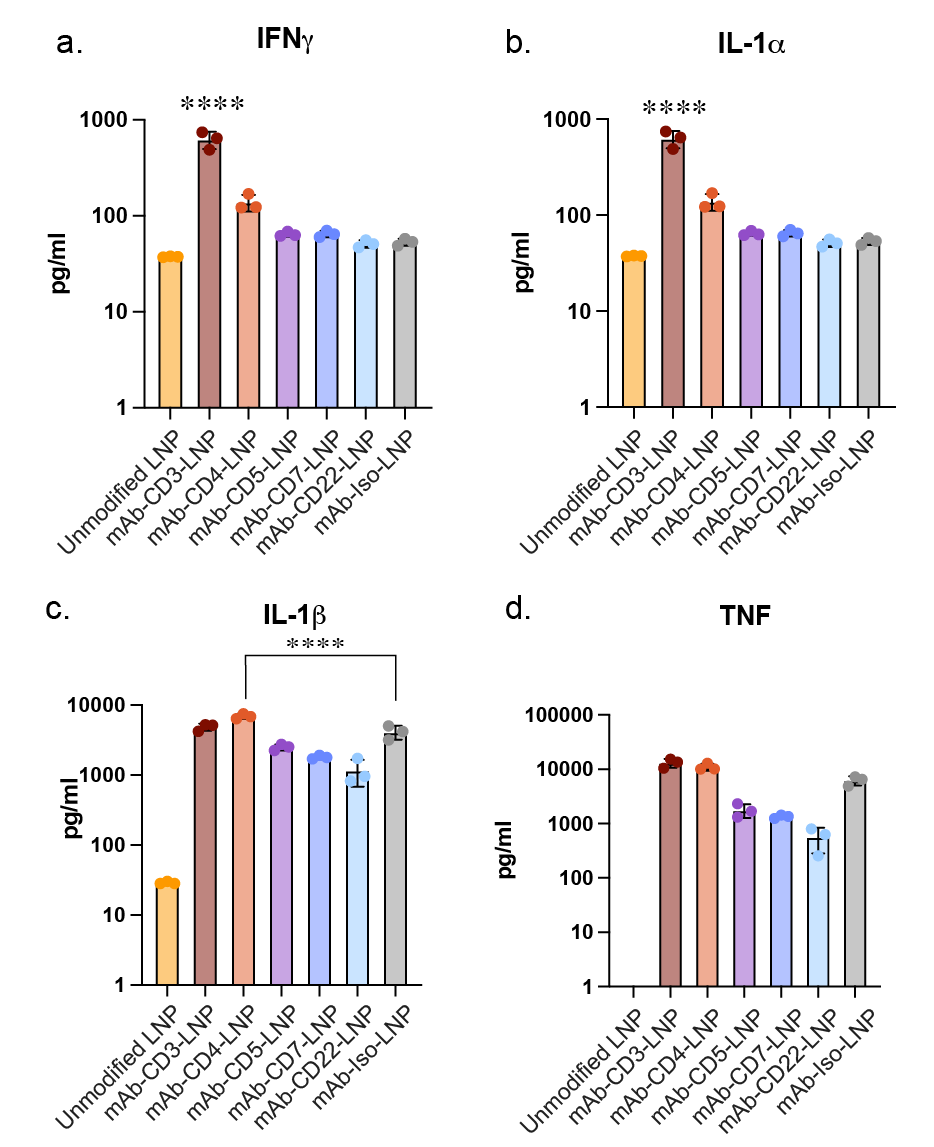

**Supplementary Figure 11. CD3 targeted LNP promoted cytokine release contributed by T lymphocytes activation.** Human whole blood was treated with 2ng/mL of different targeted LNP for 24 hours. Plasma was collected and a) IFNg, b) IL-1a, c) IL-1b and d) TNF were measured using Cytometric Bead Array (CBA) from BD biosciences following manufacture’s instruction. Detection limit (10pg/ml). ****P < 0.0001; one-way ANOVA and Dunnett's post-test. All data are means ± SD; n = 3 independent wells.

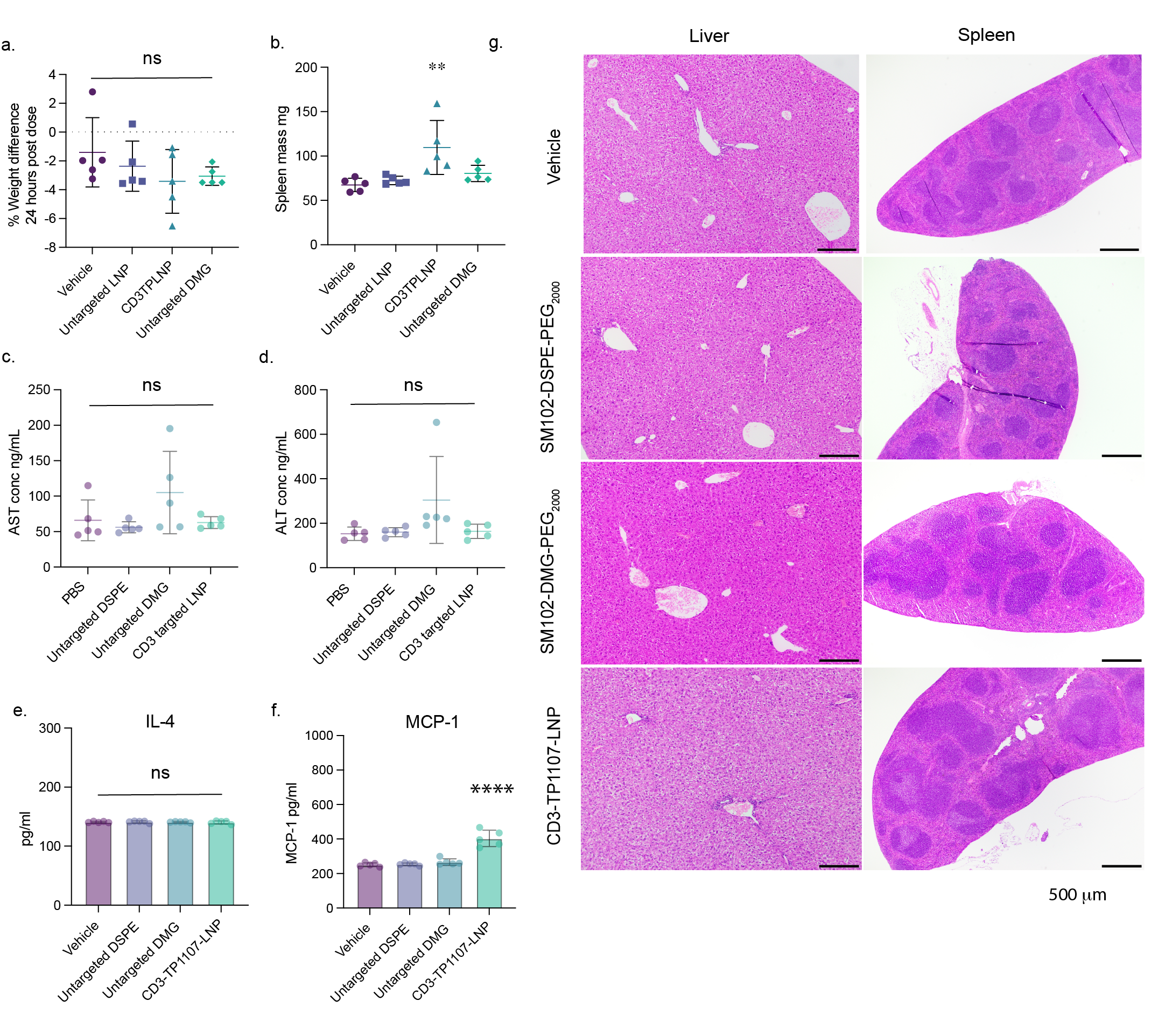

**Supplementary Figure 12. mAb-CD3-LNP did not induce obvious safety issue compared with vehicle, LNP-DSPE_2000_ and the current delivery formulation LNP-DMG_2000_.** a) Mice weight changes, b) spleen weight, c) and d) ALT/AST plasma concentration at 24 hours post dosing, e) and f) cytokines profile at 24 hours post dosing, g) histopathology examination of liver and spleen samples were measured and performed to ensure the safety aspect of the targeting system. *P < 0.05, ****P < 0.0001; one-way ANOVA and Dunnett’s multiple comparison post-test. All data are means ± SD; n = 5 individual mouse.

**Experimental Details**

**Materials**

**mRNA synthesis**

All IVT mRNA were synthesized from a PCR template containing a T7 promoter upstream followed by the codon optimized ORF. All constructs contain a 5’UTR, 3’UTR and 125 polyA tail. All mRNA was transcribed using the HiScribe T7 High Yield RNA Synthesis Kit (New England Biolabs). Capping was performed cotranscriptionally using CleanCap Reagent AG (TriLink Biotechnologies). All uridine was replaced with N 1 -Methylpseudouridine (TriLink Biotechnologies). IVT reaction was treated with DNase to eliminate the template and dsRNA was removed by cellulose clean-up methods[1]. The final product was purified by sodium acetate precipitation.

**LNP formulation**

LNPs were formulated as previously reported with modification[2]. A lipid mixture consisting of Dlin-MDA-MC3 or SM102, DSPC (Avanti Polar Lipids)), cholesterol (Sigma) and 14:0 PEG2000 PE (DMG-PEG2000) or18:0 PEG2000 PE (DSPE-PEG2000) (Avanti Polar Lipids) was prepared in ethanol as a 20 mM stock. The molar composition used was 50:10:38.5:1.5 molar ratio. The lipid solution was mixed by flowing through a micro fluidic mixing device Nanoassemblr® (Precision Nanosystems, Vancouver BC) with an aqueous mRNA solution in 10 mM citrate buffer (pH 4) at a 1:3 organic to aqueous volume ratio, 4 ml per min total flow rate. The resulting LNPs were then diluted twice with PBS (pH 7.4) immediately and further dialysed overnight. Then next day, LNPs were filtered through a 0.22 micron filter.

**LNP characterization**

The size distribution, particle number per ml and mode size of LNP was measured using Nanosight NS300 (Malvern Panalytical). The ζ potential of LNPs was measured using dynamic light scattering Zetasizer Nano ZS (Malvern Panalytical). Total mRNA content and encapsulation efficiency were determined by performing a standard Ribogreen (ThermoFisher) assay.

**Molecular cloning of sdAb.** Genes encoding the TP1107 sdAb sequence were synthesized as a gene fragment (Integrated DNA Technologies) for cloning into pET His6 TEV LIC cloning vector[3]. Plasmids will be deposited to Addgene repository.

**sdAb expression and purification.** pET-TP1107 was co-transformed alongside pEVOl-pAzF into B-95.ΔA *E. coli* which expresses the orthogonal machinery for incorporation of azPhe in recognition of UAG codon during protein translation[4]. The B-95.ΔA E. *coli* strain is a unique expression vector where 95 of its original UAG codons have been replaced along with the elimination of release factor 1 (RF-1) to facilitate improved incorporation efficiency of azPhe[5].

An overnight culture was inoculated into fresh TB media with appropriate antibiotics and grown at 37 ºC while shaking until the optical density, OD_600_ reached 0.7-1.0. sdAb expression was induced by the addition of IPTG (2 mM), L-arabinose (0.02%) and azPhe amino acid (2 mM). Protein expression was continued for a further 12-14 hours at 30 ºC before harvesting the bacteria by centrifugation. Bacterial pellets were harvested by centrifugation (4,000x g, 20 minutes) and resuspended in Ni-NTA wash buffer followed by cell lysis using a high-pressured homogeniser (Avestin Emulsiflex C5).

Upon lysis, cell debris were centrifuged (12,000x g, 30 minutes) and supernatant collected for purification via an immobilized metal affinity chromatography (IMAC) column. An additional size exclusion chromatography (SEC) was employed to remove non-specifically bound proteins using Superdex 75 10/300 GL gel filtration column (GE Healthcare). sdAb concentration was determined using Nanodrop (Thermo) spectrophotometer at 280 nm.

**Negative stain transmission electron microscopy**

Negative staining TEM was carried out by applying 3 µL of a 0.05 mg/mL solution onto a continuous carbon TEM grid (EMS 300 mesh) which has been pretreated in a plasma chamber (30 seconds, 15 mA plasma current) followed by multiple applications of Uranyl Formate (0.01 % w/v). Imaging was performed on an Thermo L120C TEM at a magnification of 92k yielding a physical pixel size of 1.55 Å/pix. 41 images were recorded on a Ceta direct electron detector. Single particle analysis was carried out in the RELION 3.1.2 software package[6]. Briefly, images had their CTF parameters estimated followed by automated particle picking and successive rounds of 2D classification to homogenize the particle stack. This yielded 19k particles for ab-initio 3D model generation, which was carried out in cryoSPARC [7] and further 3D refinements were finalized in RELION, resulting in an ~16 Å 3D reconstruction. A PDB model was generated by initially rigid body fitting of the mouse IgG (PDB: 1IGY [8]) using UCSF Chimera [9]. This fitted model then underwent an MDFF refinement using UCSF ChimeraX/iSOLDE [10, 11] to fit the antibody into the 3D volume. The resultant PDB was then used as template for HADDOCK docking [12] of the nanobody. The docking clusters that best fit the experimental density were then considered for a further round of MDFF with tight torsion and distance restraints based on the original 1IGY model and the starting model for the nanobody.

**Conjugation of sdAb to DSPE-PEG2000-DBCO**. Azide incorporated sdAbs (TP1107_optimal_) can be directly conjugated onto DBCO-PEG_2000_-DSPE through SPAAC chemistry. The conjugation mixture was prepared at molar ratio 2:1, DBCO : Azide. Meanwhile, to illustrate the effect of randomly oriented sdAbs, 2-molar excess of NHS-azide (198 Da, Thermo) was initially conjugated onto TP1107 sdAb. Excess unconjugated NHS-azide linkers were removed using a 7K MWCO Zeba desalting column (Thermo). The azide modified/incorporated sdAbs were mixed to DSPE-PEG_2000_-DBCO at a 0.5 molar excess and left for 24 hours at 37oC as TP1107_random_. No further purification was required.

**Post-insertion of active targeting module into LNPs and functionalized mAb-TP1107_optimal/random_ LNPs**

0.5% w/w of DSPE-PEG2000-TP1107 mixture was added to prepared LNPs. The reaction was incubated at 4oC for 48 hours. The free TP1107 or unreacted DSPE-PEG2000-DBCO were removed via Amicon 100 kDa MWCO (Merck) ultrafiltration. A total of 5 washes (2,000 rpm, 10 minutes) were completed. Functionalized LNP was prepared by mixing antibodies with TP1107_optimal/random_ LNP at selected ratio and incubated 4^o^C overnight.

**Calculation of TP1107 per LNP.** The sdAb number of each LNP calculated as below.

$$Number of sdAb per LNP=\frac{Number of sdAb in solution}{Number of LNP in solution}$$

Where the concentration of sdAb was calculated from western blot using JESS Simple Western™ (Bio-Techne) and the number of LNP was measured by Nanosight NS300 (Malvern Panalytical).

**Targeting antibody.** Targeting antibody that used in *in vitro* and *ex vivo* study is anti-hTfR (OKT9, purchased from WEHI facility, Victoria, Australia), mouse anti-hCD3 Antibody (UCHT1, Thermo Fisher), mouse anti-hCD4 (SK3, Biolegend), Mouse anti-hCD5 (UCHT2, Thermo Fisher), mouse anti-hCD7 (124-1D1, ThermoFisher), mouse anti-hCD22 (eBio4KB128 (4KB128), Thermo Fisher) and Mouse IgG1 kappa Isotype Control (P3.6.2.8.1, Thermo Fisher). Targeting antibody that used in *in vivo* study is mouse anti-mouse CD3ε (QA17A05, Biolegend).

**Conjugation of mAb_TfR_ to DSPE-PEG_2000_-DBCO.** A 5-molar excess of NHS-azide was initially conjugated to purified mAb_TfR_ (OKT9), a kind gift from Justine Mintern group. Excess unconjugated NHS-azide were removed by a 7K MWCO Zeba desalting column (ThermoFisher) following the manufacturer’s protocol. The azide-mAb_TfR_ were incubated with DSPE-PEG_2000_-DBCO at 37oC overnight at an DBCO : azide 2:1 ratio. No further purification is required.

**Preparation of mAb-mAbTfRLys-LNPs**

Roughly 0.05% w/w of DSPE-PEG_2000_- mAbTfR mixture was added to formulated LNPs. The reaction was incubated at 4oC for 48 hours. The post-inserted LNP was then concentrated by ultrafiltration system and slowly applied to a 90cm bed length gravity flow size exclusion column prepared with Sepharose-CL4B gel [13, 14]. The mobile phase was PBS. The fractions contained LNP were collected and concentrated by ultrafiltration system. The mRNA concentration was measured by NTA as described before. And the particle size was determined by NTA.

**Cell culture maintenance.** Jurkat cells were maintained with RPMI media (Gibco) supplied with 10% fetal bovine serum and penicillin-streptomycin (100 U/mL). Cells were cultured at 37 ºC in a humidified incubator with 5% atmospheric CO_2_ along with routine testing or mycoplasma contamination.

**Mouse models**

B6.Cg-Gt(ROSA)26Sortm14(CAG-tdTomato)Hze/J (IMSR_JAX:007914) mice were purchased from The Jackson Laboratory and maintained locally at Monash animal research platform, and mice were ordered and shipped for experiment upon requests. Male and female mice aged 6 – 14 weeks were used in experiments. All the experimental procedures followed the protocols approved by the Institutional Animal Care and Use Committee at Monash University and experimental plan was approved by the Monash Office of Research Ethics and Integrity committee under ethics 37404 or 41587. All mice group were randomized and gender balanced.

**Human PBMC collection and purification.**

Healthy donors aged between 18 – 50 years old in both sexes were recruited voluntarily after the invitation to participate. The ethics is approved by Monash University Human Research Ethics Committee, application ID 37405. The human blood was collected upon experimental plan. 10-30 mL of human blood was collected and diluted with PBS before carefully layered on Ficoll-Paque PLUS density gradient media with 1:1 v/v. PBMC layer was collected after 400g, 40 mins spin and washed with prewarmed RPMI media twice. PBMCs were either used for experiments or frozen in cell frozen media at -80oC.

**Antibody capturing LNP safety assessment and cytokine measurement**

C57BL/6 mice were acquired under ethics approval number 37404. The mice received intravenous injections of 0.1 mg/kg of various lipid nanoparticle (LNP) formulations or control groups; vehicle, unmodified SM102 with DSPE-PEG_2000_, unmodified SM102 with DMG-PEG_2000_ and mAb-CD3-LNP with SM102 and DSPE-PEG_2000_. Blood samples were collected at 6 and 24 hours post-injection to evaluate the impact on liver enzymes (ALT, AST) and the cytokine release. After 24 hours, the animals were euthanized for liver and spleen harvesting and subsequent histological analysis. The harvested liver and spleen tissues were fixed in 10% neutral buffered formalin for at least 48 hours. Two mice from each group were assessed by a veterinary pathologist for further evaluation. Plasma was obtained by centrifuging blood samples at 1,000g for 5 minutes at 4°C and stored as single aliquots. The levels of mouse cytokines IL-1α, IL-1β, IL-10, IL-6, MIP-1α, MCP-1, IL-2, TNF-α, IFN-γ, and IL-4 were measured using the BD™ Cytometric Bead Array (CBA) Mouse Flex Set according to the manufacturer's protocol. Data collection was performed using a Stratedigm S1000EXi flow cytometer.

**Cytokine measurement for the whole blood stimulated with targeted LNP**

Healthy donor’s blood was collected on the day of the experiment in heparin-coated collection tubes. Different targeted LNP were added to each well at a final concentration of 2 ng/μL and incubated for 24 hours. The blood samples were then centrifuged to collect plasma for cytokine measurement. Human cytokines IFN-γ, IL-1α, IL-1β, and TNF were quantified using the BD™ Cytometric Bead Array (CBA) Human Flex Set according to the manufacturer's protocol. Data collection was performed using a Stratedigm S1000EXi flow cytometer.

**Cells association and transfection assay with functionalized LNPs.** To assess the binding and transfection efficiency of functionalized lipid nanoparticles (LNPs) in Jurkat cells, approximately 50,000 or 100,000 cells were added to individual wells in a 96-well plate. A final concentration of 0.5 ng/µL or 1 ng/µL of mRNA was added to the cells and incubated at 37°C for varied time periods. Subsequently, cells were washed three times with 2% FBS-PBS following centrifugation at 400x g for 5 minutes. Cells were then resuspended in 80 µL of 2% FBS-PBS, and the mean fluorescence intensity was quantified using a Stratedigm S1000EXi flow cytometer. eGFP and Cy5 fluorescence were excited at 488 and 642 nm, respectively, with fluorescence emission collected at 520/20 nm and 676/29 nm.

**Human PBMC association and transfection assay with functionalized LNPs.** To assess the binding and transfection efficiency of the functionalized LNPs approximately 500,000 PBMC were added to individual wells in a 96-well plate with functionalized LNPs at 1ng/µL final concentration. Then cells were incubated at 37^o^C for 24 hours. PBMC then were washed thrice with 2% FBS-PBS after centrifugation at 400x g for 5 minutes. To phenotype the sub-populations, cells were stained against αCD3-PE mAb (clone OKT3, Biolegend), αCD4-BV510 mAb (clone OKT4, Biolegend), αCD8-BV786 mAb (clone SK1, Biolegend), αCD19-BV421 mAb (clone HIB19, Biolegend), αCD14-Alexa Fluor 700 mAb (clone HCD14, Biolegend), αCD56-BV605 mAb (clone 5.1H11, Biolegend), and viability dye (eBioscience™ Fixable Viability Dye eFluor™ 780, Thermofisher) on ice for 30 min. Antibodies were all used at 1:200 dilutions with Human TruStain FcX™ (Biolegend) as manufacturer’s protocol. After washing away the excessive antibody, cells were resuspended with 100 µL 2% FBS-PBS for the flow analysis (Stratedigm S1000EXi). Cells were identified by a combination of surface markers: CD4+ T cells (CD3+ and CD4+), CD8+ T cells (CD3+ and CD8+), monocytes (CD3-, CD19-, CD56-, and CD14+), NK cells (CD3-, CD19-, CD14- and CD56+), and B cells (CD3− and CD19+). eGFP and Cy5 fluorescence was excited at 488 and 642 nm with fluorescence emission collected at 520/20 nm and 676/29 nm respectively. The data in supplementary figure 8 was obtained via Cytek Aurora 5 lasers full spectrum cytometer.

**In vivo assessment of CD3 targeting of LNPs to T cells cross multiple organs**

Ai14 mice were injected intravenously with unmodified lipid nanoparticles (LNPs), CD3-targeted LNPs, or isotype control LNPs loaded with Cre mRNA. After 24 hours, blood was collected via cardiac puncture, and the mice underwent transcardiac perfusion with PBS to remove circulating blood. Red blood cells were lysed using ACK buffer (Thermo Fisher, USA) at a 1:10 (v/v) ratio twice, followed by washing with 2% FBS-PBS. The liver, spleen, and lymph nodes (inguinal, iliac, and cervical) were collected and processed as follows:

The liver was minced and digested using a gentleMACS™ dissociator with 2.8 mg/mL Collagenase H and 0.28 mg/mL DNase. The digested mixture was filtered to remove undigested material and subjected to a slow spin at 60g. The supernatant was collected and spun down to collect the pellet. The pellet was resuspended in 30% Percoll media and spun to remove hepatocytes, followed by resuspension with ACK lysis buffer and washing with 2% FBS HBSS before antibody staining.

The spleen was minced with 1 mg/mL Collagenase III and 0.28 mg/mL DNase and digested by constant gentle mixing until fully digested. Cells were filtered and red blood cells lysed using ACK buffer.

Lymph nodes were collected and homogenized by passing through a 0.45 µm filter. Dissociated cells were collected and washed with media.

All immune cell pellets were stained with a flow cytometry panel containing the following antibodies: αCD3e-BV650 mAb (clone 145-2C11, BD Biosciences), αCD90.2-BV650 mAb (clone 53-2.1, BD Biosciences), αCD4-APC-Cy7 mAb (clone GK1.5, BioLegend), αCD8-BV711 mAb (clone 53-6.7, BD Biosciences), αCD19-BV786 mAb (clone 1D3, BD Biosciences), αCD11b-BV421 mAb (clone M1/70, BioLegend), αLy-6C-BUV661 mAb (clone HK1.4.rMAb, BD Biosciences), αLy-6G-BV605 mAb (clone 1A8, BD Biosciences), αCD45-Pacific Blue mAb (clone S18009F, BioLegend), αI-A/I-E-BV510 mAb (clone M5/114.15.2, BioLegend), αF4/80-PE/Dazzle mAb (clone BM8, BioLegend), and αCD11c-Alexa Fluor 700 mAb (clone N418, BioLegend). Additionally, Mouse BD Fc Block™ and viability dye (LIVE/DEAD™ Fixable Blue Dead Cell Stain Kit, Thermo Fisher) were included. Samples were incubated on ice for 30 minutes, followed by washing to remove excess antibody.

Flow cytometry was performed using a Cytek Aurora 5 laser cytometer, and data were analyzed using FlowJo (BD Biosciences). Leukocyte phenotyping was conducted using the following markers: CD4+ T cells (CD45+, CD11b-, CD3e+, CD4+); CD8+ T cells (CD45+, CD11b-, CD3e+, CD8+); dendritic cells (CD45+, CD3e-, CD19-, CD11c+, MHCII+); monocytes (CD45+, CD11b+, Ly6C+, Ly6G-); neutrophils (CD45+ CD11b+, Ly6C+, Ly6G+); macrophages (CD45+ CD11b+, Ly6C low, Ly6G-, F4/80+, SSA low); and CD19+ B cells (CD45+, CD11b-, CD3-, CD19+ or MHCII+).

**Statistical analysis.** Data are presented as mean ± standard deviation based on the data obtained from at least n = 3 independent experiments or well or mice. Statistical significance was determined using GraphPad Prism 9.0 and stated in each figure legend.

**References:**

1.	Baiersdörfer, M., et al., *A facile method for the removal of dsRNA contaminant from in vitro-transcribed mRNA.* Molecular Therapy-Nucleic Acids, 2019. **15**: p. 26-35.

2.	Veiga, N., et al., *Cell specific delivery of modified mRNA expressing therapeutic proteins to leukocytes.* Nat Commun, 2018. **9**(1): p. 4493.

3.	Pleiner, T., M. Bates, and D. Gorlich, *A toolbox of anti-mouse and anti-rabbit IgG secondary nanobodies.* J Cell Biol, 2018. **217**(3): p. 1143-1154.

4.	Chin, J.W., et al., *Addition of p-Azido-l-phenylalanine to the Genetic Code of Escherichia c oli.* Journal of the American Chemical Society, 2002. **124**(31): p. 9026-9027.

5.	Mukai, T., et al., *Highly reproductive Escherichia coli cells with no specific assignment to the UAG codon.* Scientific Reports, 2015. **5**(1): p. 9699.

6.	Zivanov, J., et al., *New tools for automated high-resolution cryo-EM structure determination in RELION-3.* elife, 2018. **7**: p. e42166.

7.	Punjani, A., et al., *cryoSPARC: algorithms for rapid unsupervised cryo-EM structure determination.* Nature methods, 2017. **14**(3): p. 290-296.

8.	Harris, L.J., E. Skaletsky, and A. McPherson, *Crystallographic structure of an intact IgG1 monoclonal antibody.* J Mol Biol, 1998. **275**(5): p. 861-72.

9.	Goddard, T.D., C.C. Huang, and T.E. Ferrin, *Visualizing density maps with UCSF Chimera.* J Struct Biol, 2007. **157**(1): p. 281-7.

10.	Meng, E.C., et al., *UCSF ChimeraX: Tools for structure building and analysis.* Protein Sci, 2023. **32**(11): p. e4792.

11.	Croll, T.I., *ISOLDE: a physically realistic environment for model building into low-resolution electron-density maps.* Acta Crystallogr D Struct Biol, 2018. **74**(Pt 6): p. 519-530.

12.	Honorato, R.V., et al., *Structural Biology in the Clouds: The WeNMR-EOSC Ecosystem.* Front Mol Biosci, 2021. **8**: p. 729513.

13.	Breda, L., et al., *In vivo hematopoietic stem cell modification by mRNA delivery.* Science, 2023. **381**(6656): p. 436-443.

14.	Rurik, J.G., et al., *CAR T cells produced in vivo to treat cardiac injury.* Science, 2022. **375**(6576): p. 91-96.
